# Single component CRISPR-mediated base- editors for *Agrobacterium* and their use to develop an improved suite of strains

**DOI:** 10.1101/2024.08.04.606528

**Authors:** Vincent J. Pennetti, Peter R. LaFayette, Wayne Allen Parrott

## Abstract

Agrobacterium mediated plant transformation largely depends on two distinct strain lineages – C58 and Ach5. To better serve the plant transformation community, we have created a suite of auxotrophic and auxotrophic recombinant deficient mutants of C58 derivatives EHA105, GV3101::pMP90, and Ach5 derivative LBA4404. While these derivatives are useful, having additional strain backgrounds available would help expand the repertoire for plant transformation even further. Toward that end, two underutilized hypervirulent strains are K599 (NCPPB 2659), and Chry5—but disarmed variants are not easily accessible. To improve availability, we produced disarmed versions of *A. rhizogenes* strain K599 and *A. tumefaciens* strain Chry5 and introduced the same desirable mutations as with the other lineages. Each thymidine auxotrophy and recombination deficiency were introduced to existing and newly disarmed Agrobacterium strains via loss of function mutations conferred to *thyA* and *recA*, respectively, through CRISPR-mediated base-editing of codons amenable to nonsense mutation. To streamline the editing process, we created a series of visually marked single component base-editor vectors and a corresponding guide-filtering Geneious Prime wrapper plugin for expedited guide filtering. These new strains, the simplified CRISPR-mediated base-editor plasmids, and streamlined workflow will improve the ease with which future *Agrobacterium* strain derivatives are created while also supporting plant transformation at large.

## Introduction

The creation of special-use strains of *E. coli* have facilitated developments in molecular biology. A variety of enhancing modifications have been introduced to increase plasmid stability, transformation efficiency, and enable blue-white screening [1].

*Agrobacteria* are to plant transformation what *E. coli* is to molecular biology, but in contrast to *E. coli*, little has been done to specialize available *Agrobacterium* mutants for plant transformation.

*Agrobacterium*-mediated transformation is currently the most widely adopted method for plant transformation, supporting both public and private sector efforts at enhancing food security. *Agrobacterium* is the causal agent of two unique disease phenotypes, crown gall disease and hairy root disease, caused by *Agrobacterium tumefaciens* and *Agrobacterium rhizogenes* , respectfully [2]. Exploiting the natural process of DNA transfer by replacing the disease-causing genes with genes of interest is the basis of using *Agrobacterium* as a transformation tool.

Phytopathogenic strains of *Agrobacterium* possess a virulence plasmid encoding four main functions: plant oncogenicity and opine anabolism, virulence, opine catabolism, and regulatory/conjugation/replication [3]. The oncogenic and opine anabolism genes for crown gall disease and hairy root disease are located a large plasmid in the bacterium known as the Ti/Ri, or “tumor-inducing”/ “root-inducing”, plasmid. The oncogenic genes themselves are located within 25-bp “border” sequences which delineate the sequence of DNA that ultimately is transferred to the plant [4].

Eliminating the bacterium’s ability to cause disease while maintaining virulence through interruption of the oncogenic genes, or “disarming”, has transformed *Agrobacterium* into an indispensable tool for genetic engineering.

While there are many named strains for plant transformation, these represent only two lineages of disarmed *Agrobacterium* derived from C58 and Ach5 [5](Figure 1). The C58 lineage includes strains GV3101::pMP90 [6], EHA101 [7], EHA105 [8] while LBA4404 [9] is the only popularized disarmed strain from the Ach5 lineage. Aside from being disarmed, all these strains are essentially wildtype in the rest of their genome. The only popularized exception is AGL-1 [10], a C58 derivative with recombination deficiency conferred by a deletion of the *recA* locus. Other reports of *recA* minus strains have been impractical to use as they lacked virulence functions [11]. One notable exception to the lack of beneficial mutations *Agrobacterium* strains intended for plant transformation can be seen through thymidine auxotrophic LBA4404 [12, 13] which has improved biocontainment and elimination properties.

**Figure 1.**
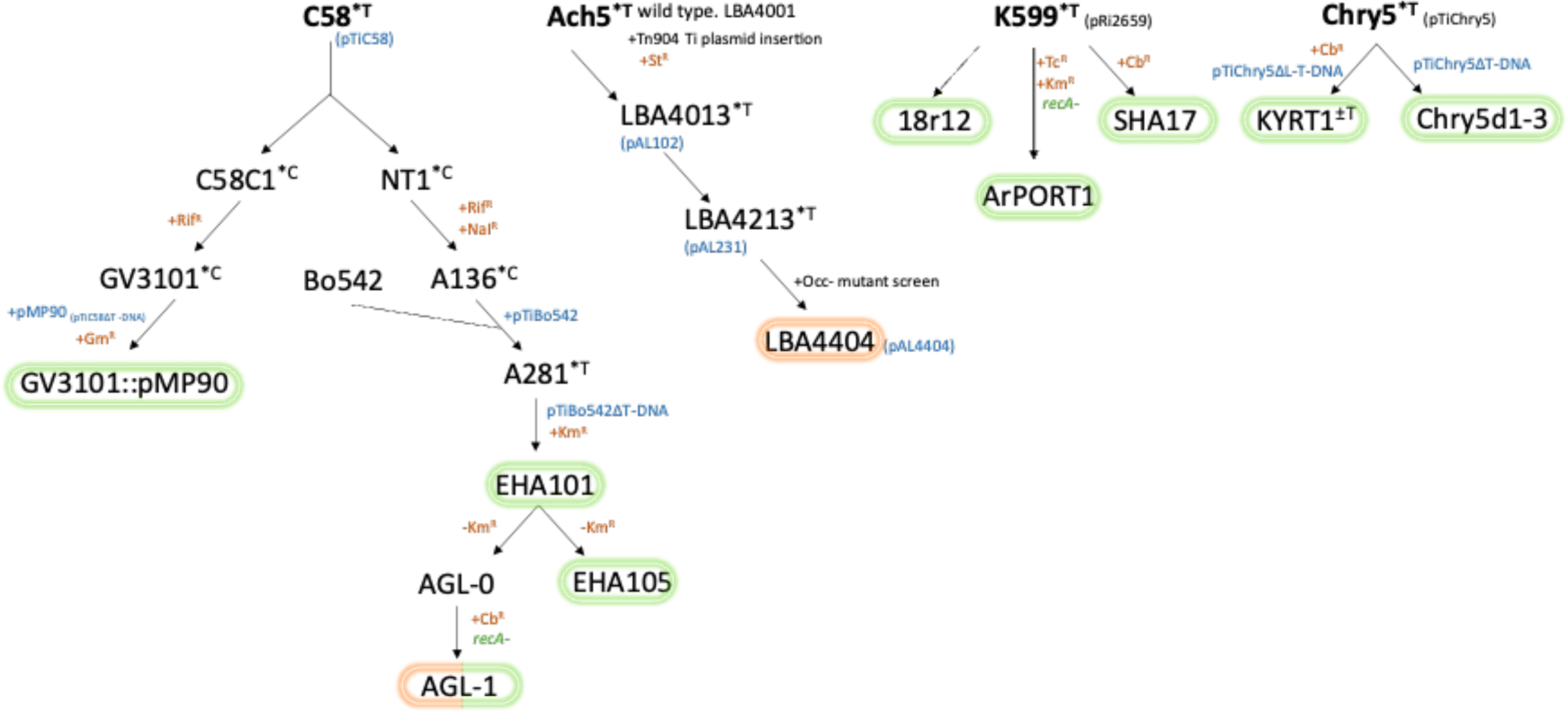
Pedigree of current laboratory strains of *Agrobacterium* as modified from De Saeger et al. (2021). Circled strains are common laboratory strains. Strains circled in orange were disarmed through transposon mutagenesis, strains in green were disarmed through homologous recombination. Note that of these strains, 18r12, SHA17 and KYRT1 are not readily available and hence are used in a minority of transformations. ArPORT1 was specifically made as cointegrate acceptor for GA*A*NTRY-based cloning, and contains a C58 right border on the disarmed Ri plasmid [44]. Further, Chry5d1, Chry5d2, and Chry5d3 are all privately held by Syngenta. *T armed Ti plasmid; *C cured of Ti plasmid, +Cb^R^ introduced carbenicillin resistance, +Gm^R^ introduced gentamycin resistance, +Km^R^ introduced kanamycin resistance, +Nal^R^ introduced nalidixic acid resistance, +Rif^R^ introduced rifampicin resistance, +St^R^ introduced streptomycin resistance, +Tc^R^ introduced tetracycline resistance.

Molecular tools are being developed faster than *Agrobacterium* strains. Historically, *Agrobacterium* mutants were generated through multiple strategies including UV mutagenesis [14], transposon mutagenesis [15], and homologous recombination [7]. These technologies were relied upon due to a lack of sequencing information. CRISPR- Cas9 has become an indispensable tool for eukaryotic genome engineering through double stranded DNA, or DSB, break-repair mediated gene knockouts now that DNA sequencing is more accessible. Gene knockouts rely on imperfect DSB repair mechanisms such as non-homologous end joining, or NHEJ [16]. While DSB repair is efficient for gene knockout in eukaryotes, *Agrobacterium* lacks the NHEJ pathway.

Therefore, when DSBs are made in *Agrobacteria*, it destabilizes the genome and often results in lethality [17].

CRISPR-derived systems that do not induce double stranded breaks, such as base-editing, circumvent the lethality of DSBs. Base-editing with the Target-AID base editing architecture [18] composed of a nuclease-dead Cas9, or dCas9, and a cytidine deaminase, such as *Pm*CDA1, efficiently generate premature stop codons by converting the cytidine residues to thymidine within 5’-CAA-3’, 5’-CAG-3’, 5’-CGA- 3’, or 3’-CCA-5’ codons without causing a DSB. Use of the Target-AID architecture is highly efficient in *A. tumefaciens* and *A. rhizogenes* [19], as well as other phytopathogenic bacteria like *Pseudomonas* [20]. Editing with the Target-AID architecture has enabled rapid mutant generation within existing *Agrobacterium* lineages [19, 20].

There are three main challenges when adopting a system for generating base-edited mutants of *Agrobacterium*. The first challenge is that current base-editors for phytopathogenic bacteria rely upon Gateway assembly to produce the final editor plasmid [19, 20]. Gateway assembly is expensive to implement and largely unnecessary for the deployment of the editors. Gateway assembly is only present in the current editor workflow to stack the four required elements for editing in this fashion: a backbone containing a compatible replicon for the host, *sacB* for plasmid eviction, a sgRNA expression cassette, and the base-editor itself. The multi-component modularity was essential for rapid development of the editor plasmids when specific gene combinations were being evaluated for efficiency, but subsequently became an unnecessary barrier to deployment at scale now that they have characterized on target and spurious editing efficiencies.

The second challenge with current generation base-editors for phytopathogenic bacteria is plasmid eviction. The *sacB* marker is used in conventional microbe engineering as it is an effective counter selective agent to screen for isolates that have evicted the plasmid bearing it when plated on a sucrose-containing medium. However, a fraction of colonies plated on sucrose still fail to evict the plasmid on first plating. Li et al. (2023) addressed this issue by delivering GFP in tandem with the final editor replicon.

However, GFP requires additional hardware for visualization, and their editor plasmid still relies on Gateway assembly each time a new gRNA is to be assembled. Further, sfGFP is already used earlier in the assembly process to monitor gRNA cloning [19], rendering a two-step restriction ligation and Gateway assembly unavoidable to take advantage of the visual reporters with overlapping spectra [20].

A third challenge with current-generation base-editors lies in efficient guide filtering. While CRISPR-mediated base-editors like Target-AID rely upon dCas9 to bind DNA, not all binding sites compatible with dCas9 are amenable for cytidine-deaminase mediated gene knockout. Current prediction algorithms tailored to conventional Cas9- mediated knockouts output far more putative guides than can be used for base-editing mediated loss of function [21, 22]. One strategy to refine the input requirements for gRNA generations so that only sequences containing cytidine residues within the editing window of a predicted guide are produced. While this dramatically cuts down on the number of guides generated, there remains a large output of putative guides that need to be curated with respect to the reading frame, a process that can be tedious and time-consuming.

Command line and web-based tools like CRISPR-CBEI can expedite the guide filtering process [23]. CRISPR-CBEI provides the option to predict guides and their off-target effects for a variety of different base-editor architectures. CRISPR-CBEI, however, requires either upload of self-generated genome files for off-target analysis on an externally hosted platform, or interaction through the command line to perform the analysis locally. Many researchers design their constructs *in silico* prior to producing them in the lab in software such as SnapGene or Geneious Prime. Using CRISPR-CBEI either in its web interface or through the command line requires users to bypass their plasmid design software to curate guide selection and then later manually annotate their digital maps before cloning, further exacerbating the inefficiencies of current base-editing workflows.

Here, we sought to address the three main inefficiencies of current base-editing workflows for *Agrobacterium* strain engineering: by 1) constructing a suite of single-component base editor vectors for streamlined cloning, 2) introducing chromoproteins that can be visualized without additional hardware to monitor plasmid eviction, and 3) developing a Geneious Prime wrapper plugin to simplify the guide prediction and annotation for *Agrobacterium* mutant generation. Using the simplified vectors and workflow, we generated a suite of derivative strains with desirable mutations conferring thymidine auxotrophy and recombination deficiency, to expand the utility of current laboratory strains of *Agrobacterium*. Further, we also produced disarmed derivatives of the hypervirulent *A. rhizogenes* K599 and the difficult-to disarm *A. tumefaciens* Chry5 using different allelic replacement strategies, and subsequently introduced the same desirable mutations to the newly disarmed variants. The two desirable mutations introduced into each background were produced serially, and off-target effects of this strategy were analyzed through whole genome sequencing. Our single component base-editor vectors will be made available through Addgene, and the Geneious Prime wrapper plugin is currently available through GitHub for adoption or modification.

## Materials and Methods

### Molecular biology reagents and methods

All plasmids were produced using NEB (New England Biolabs, Ipswich, MA) restriction ligation and Gibson assembly reagents including *Bsa*I-HF-v2 (NEB #R3733S), NEBuilder HiFi DNA Assembly Master Mix (NEB #E2621S) and *E. coli* cloning strain 10-beta (NEB # C3019H). *E. coli* strain One Shot *ccd*B Survival cells (ThermoFisher Scientific, Waltham, MA, #A10460) was used for cloning and plasmid propagation of all materials containing the *ccd*B/CmR gene cassette.

Oligos and primers used for Gibson assembly were synthesized by Millipore Sigma (Millipore Sigma, Burlington, MA) and can be found in Supplementary Table 1. PCR products were produced using Q5 high fidelity polymerase (NEB # M0491S) under the recommended three-step amplification conditions unless otherwise stated. All NEBuilder HiFi assemblies were executed following standard incubation [24] and transformation procedures [25] into their respective destination strains. All plasmids were isolated using standard protocols and reagents from the GenCatch plasmid DNA mini-prep kit (Epoch Life Science, Missouri City, TX). Plasmids produced using amplified products were sequenced with Oxford Nanopore long-read sequencing via Plasmidsaurus (Plasmidsaurus, Eugene, OR). Successfully cloned sgRNA targets were validated with Sanger sequencing (Genewiz, South Plainfield, NJ) using oligo 5- GTATGATGAGAGCACCGATGAG-3’. A list of assemblies used in this study can be found in Supplementary Table 2.

### Strains, media, and culture conditions

For of *E. coli* transformation, 2 µL of assembly mix were combined with 10 µL of freshly thawed competent cells, incubated on ice for 30 minutes in a 2-mL microcentrifuge tube, heat shocked at 42°C for 30s and immediately returned to ice for 5 minutes. Two hundred µL of resuspension medium (SOC for DH5α and OneShot cells, NEB 10-beta stable outgrowth medium for 10-beta cells) was added to the tubes, and the bacterial suspensions were then incubated for 1 hr at 37°C prior to plating on LB with appropriate antibiotics (50 µg·mL^-1^ kanamycin, 100 µg·mL^-1^ spectinomycin, 10 µg·mL^-1^ tetracycline). Colonies were isolated for liquid culture and sequencing following overnight incubation at 37°C.

*Agrobacterium* electrocompetent cells were prepared and used as specified in the BioRad MicroPulser™ (Biorad Laboratories, Hercules CA, US) manual unless otherwise noted. Thymidine was added to the recovery medium for any strain auxotrophic for thymidine or receiving an editor plasmid for thymidine auxotrophy. YP medium [26] was used in lieu of LB medium for all strains rendered *recA* deficient or as the recovery medium for strains receiving a *recA* editor plasmid. All *Agrobacterium* derivatives are listed in their full genotype and corresponding shorthand nomenclature of the form: “PEN” for Pennetti; followed by a unique number. Generally, the first digit of the number designates a lineage, the second digit a modifier—0 for disarmed, 2 for armed, the third digit a mutation. All strain derivatives are listed in Table 1.

**Table 1.**
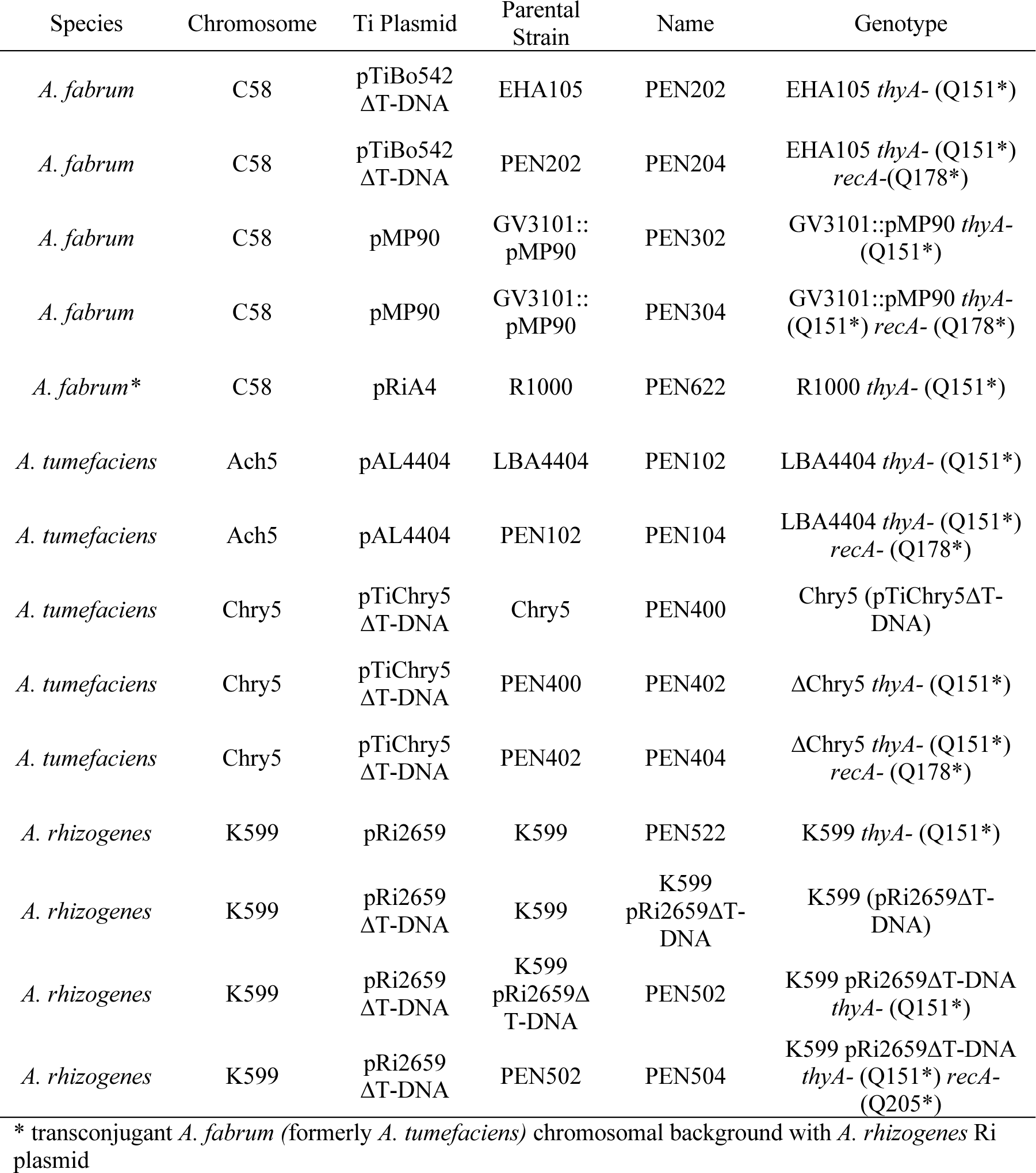
List of base edited strains produced with the single component base-editors. The respective species, chromosomal background Ti Plasmid, parental strain, name of the derivative, and corresponding genotype are listed.

### Single component base-editor vector construction

The first single component base-editor vector was initially produced by reconstituting the Gateway vector originally proposed for base editing in *Agrobacterium* [19] using NEBuilder assembly. Single component base editor pSpCas9d-CDA1-sfGFP-SacB- GGPK/pVP141 was produced through amplification of three fragments from the original Gateway plasmids[19] using overlapping oligos and a *Bsa*I-HFv2 restriction digest of pGGPK-AG2 as the recipient backbone. Fragments containing the Target-AID expression cassette, the sfGFP excision marker for guide assembly and scaffold sequence, and the *sacB* eviction marker were amplified from each pEN-L4-PvirB- dCas9-CDA-UL-T3T-R1, pEN-L1-PJ23119-Bsa1-PglpT-sfGFP-TrrfB-Bsa1-Scaf-L2, and pEN-R2-SacB-L3, respectively [19]. The amplicon containing the editor cassette was amplified using Phusion Plus Green PCR Master Mix (ThermoFisher Scientific, Waltham, MA, #F632S) using the recommended three-step amplification conditions, while the other amplicons were produced using Q5 high fidelity polymerase (NEB # M0491S). Primers used for amplification are in Supplementary Table 1. Amplicons were isolated from a 0.5X TBE gel by centrifugation of a gel core through EconoSpin column (Epoch Life Science, Missouri City, TX), at 10000 x *g* for three minutes. One µL of each eluate was then combined with 1 µL of *Bsa*I-HFv2 digested pGGPK-AG2 normalized to 50 ng·µL^-1^ and 2.5 µL of NEBuilder HiFi DNA Assembly Master Mix and incubated at 50°C for 1 hour prior to transformation into chemically competent *E. coli* 10-beta cells. Colonies were visually screened for sfGFP expression prior to sequencing.

To expand the utility of the single component base-editor vector, a series of chromoprotein-marked *sacB* containing vector backbones were produced prior to combining them with the editor machinery. The eforRed chromoprotein [27] was synthesized as a gBlock fragment (IDT, Newark, NJ) with the bacterial PJ23100 promoter and B0030 RBS. PCR amplicons were generated using Q5 high fidelity polymerase with overlapping primers listed in Supplementary Table 1, corresponding to eforRed, a *ccd*B/CmR cassette amplified from pGGP-AG, and a *sacB* expression cassette amplified from pSpCas9d-CDA1-sfGFP-SacB-GGPK/pVP141. These amplicons were assembled with a PCR amplicon from either the pGGP-AG or pGGPK- AG2 backbones without *Agrobacterium* border repeats using 2.5 µL of NEBuilder HiFi DNA Assembly Master Mix and incubated at 50°C for 1 hour prior to transformation into chemically competent One Shot *ccd*B Survival cells, to generate pVPHDB- Se/pVP073 and pVPHDB-Ke/pVP074, respectively. Once verified, pVPHDB- Ke/pVP074 was used as template for inverse PCR to exchange eforRed’s coding sequence with that for the AeBlue chromoprotein from *Actinia equina* [27], generating pVPHDB-KAe/pVP075.

Chromoprotein-marked, single component base-editor vectors lacking unnecessary T-DNA border repeats were then generated using each pVPHDB-Se/pVP073 and pVPHDB-Ke/pVP074 as the backbone. Independent 10-µL restriction digests were set up for pVPHDB-Se/pVP073 and pVPHDB-Ke/pVP074 using 1 µg of plasmid, 0.5 µL of each *Hind*III-HF (NEB #R3104S) and *Eco*RV-HF (NEB #R3195S), and 1 µL of CutSmart Buffer (NEB #B6004S) and incubated for one hour at 37°C prior to a 20- minute heat inactivation at 80°C. A PCR amplicon with 24-27 bp of complementarity on either end to the digested vectors was produced from pSpCas9d-CDA1-sfGFP- SacB-GGPK/pVP141 encompassing the expression cassettes for the Target-AID editor, the sfGFP excision marker, and for the scaffold. The amplicon was isolated through a column as before. One µL of the eluate was combined with 1 µL of each heat-inactivated digest (pVPHDB-Se/pVP073 and pVPHDB-Ke/pVP075) and assembled with 1 µL of NEBuilder HiFi DNA Assembly Master Mix. The assemblies were transformed into chemically competent *E. coli* 10-beta cells and selected on LB medium containing each spectinomycin and kanamycin generating pSpCas9d-CDA1- sfGFP-SacB-KAe /pVP143 and pSpCas9d-CDA1-sfGFP-SacB-Se/pVP144, respectively. Correctly assembled colonies were also screened for their respective visual reporters prior to whole plasmid sequencing via Plasmidsaurus.

### Guide prediction

Guide prediction for all targets was performed through Geneious Prime using a Geneious Prime wrapper plugin connected to a simple Python script originally named the Base Editor Enrichment Function, or BEEF.

### Base-edited strain generation

Single component base-editors were used in generating a suite of thymidine-auxotrophic and *recA* minus *Agrobacterium strains*. Single stranded oligos for the respective sgRNA targets are listed in Supplementary Table 1. Each oligo was designed to have19 bp of complementarity to each overhang left behind by *Bsa*I-HFv2 digestion of the single component base editor plasmids. For guide insertion, 1 µg of plasmid DNA was digested using 0.5 µL of *Bsa*I-HFv2 in a 10 µL reaction with 1 µL of CutSmart buffer for 1h and heat inactivated at 80°C for twenty minutes. A 1:250 dilution of the 100 µM oligos (Millipore Sigma, Burlington, MA) were prepared with type I water. One µL of each oligo dilution was combined with 1 µL of the desired *Bsa*I-HFv2 digested editor plasmid and 1 µL of NEBuilder HiFi DNA Assembly Master Mix and incubated at 50°C for 1 hour prior to transformation into chemically competent *E. coli* 10-beta cells. Successful assemblies were scored for lack of sfGFP expression prior to Sanger sequencing.

Once a correct assembly was identified, electrocompetent cells of the recipient *Agrobacterium* strain were prepared as specified in the BioRad MicroPulser™ (Biorad Laboratories, Hercules CA, US) and the plasmid was electroporated. Electroporations targeting thymidine auxotrophy were recovered in liquid LB medium supplemented with 150 mg·L^-1^ thymidine and no antibiotics for 2 h at 28°C with shaking, while electroporations targeting *recA* were recovered in liquid YP medium supplemented with 150 mg·L^-1^ thymidine and no antibiotics for 4 h at 28°C with shaking. Two hundred µL of the recovery medium was streaked on either solid LB or YP medium supplemented with appropriate antibiotics and incubated at 28°C. Once colonies were visible, single colonies were restreaked on the same medium for additional incubations at 28°C to recover edited lines. The resulting colonies were then restreaked onto LB or YP medium containing 10% sucrose for plasmid eviction. Colonies failing to express the chromoprotein reporter on the plasmid backbone were screened via PCR spanning the desired edit, and amplicons were run on a gel, purified with the Zymo gel DNA recovery kit (Zymo research, Orange, CA, #D4001), and sent for Sanger sequencing. Editing outcomes were verified using EditR (https://github.com/MoriarityLab/EditR). Colonies containing edits were subsequently restreaked on sucrose-containing medium and sequenced a second time before freezing stock cultures. Thymidine auxotrophic targets were phenotypically validated via replicate plating with and without exogenous thymidine supplementation prior to Sanger sequencing.

Thymidine auxotrophs of EHA105 and LBA4404 were produced using pSpCas9d- CDA1-sfGFP-SacB-GGPK/pVP141-derived editor plasmids. Chry5 base-edited strains were produced using pSpCas9d-CDA1-sfGFP-SacB-Se/pVP144-derived editor plasmids. All remaining base-edited strains isolated were produced using pSpCas9d- CDA1-sfGFP-SacB-KAe /pVP143-derived editor plasmids. All edited strain derivatives produced using the single component editor plasmids are described in Table 1. *RecA* mutants were produced serially, after thymidine auxotrophy was confirmed in the respective background.

### Single component base editor eviction

Single component base-editors pSpCas9d-CDA1-sfGFP-SacB-KAe /pVP143 and pSpCas9d-CDA1-sfGFP-SacB-Se/pVP144 carrying sgRNAs targeting the *thyA* gene in each EHA105, K599 pRi2659ΔT-DNA, LBA4404, and PEN400 were cloned using single-stranded oligos as before. Both pSpCas9d-CDA1-sfGFP-SacB-KAe /pVP143 and pSpCas9d-CDA1-sfGFP-SacB-Se/pVP144 derived editors were made for each background, except PEN400, for which only a pSpCas9d-CDA1-sfGFP-SacB- Se/pVP144 derived editors were cloned. The plasmids were electroporated into their recipient strains and plated on YP medium supplemented with 150 µg·mL^-1^ thymidine and appropriate antibiotics for the plasmid before incubating until single colonies emerged. Once single colonies could be identified, three colonies from each electroporation were inoculated into 3mL YP cultures supplemented with 150 µg·mL^-1^ thymidine and appropriate antibiotics for overnight incubation at 28°C with agitation at 240 rpm. The following day, the OD600 of each culture was normalized to 0.1 and serial dilutions were plated through 10^-5^ onto YP medium containing 10% sucrose for editor plasmid eviction. Resulting colonies on the serial dilution plates with sufficient separation were scored for presence/absence of the chromoprotein reporter in each colony that formed. Data from each triplicated strain was imported to GraphPad Prism10 for macOS version 10.2.0 for plotting and analysis.

### MMS screen

Base-edited *recA* mutants were evaluated for mutagen sensitivity along with their progenitor strains [19]. EHA105, PEN202, PEN204, LBA4404, PEN102, PEN104, K599 pRi2659ΔT-DNA, PEN502, PEN504, PEN400, PEN402, and PEN404 were inoculated into 3-mL YP cultures supplemented with 150 µg·mL^-1^ thymidine for overnight incubation at 28°C with agitation at 240 rpm. The following day, six serial dilutions were made from each culture ranging from OD600 10^-1^ to 10^-6^ in a 96-well plate format. A 3-µL drop from each serial dilution was spotted on a solid YP 150 µg·mL^-1^ thymidine plate supplemented with or without 0.005% (v/v) methyl methanesulfonate (MMS) and incubated at 28°C for 48 h.

### Growth curves

Growth performance for each lineage except LBA4404 was evaluated by incubation and spectrophotometric quantification in a Biotek Synergy2 plate reader (BioTek Instruments Inc., Winooski, VT, USA). As with the MMS screen, EHA105, PEN202, PEN204, K599 pRi2659ΔT-DNA, PEN502, PEN504, PEN400, PEN402, and PEN404 were inoculated into 3-mL YP cultures supplemented with 150 µg·mL^-1^ thymidine for overnight incubation at 28°C with agitation at 240 rpm. The following day, 2 mL of each liquid culture was pelleted at 8000 x *g* and washed 1x with PBS (137 mM NaCl, 2.7 mM KCl, 10 mM Na2HPO4, 1.8 mM KH2PO4). After PBS washing, cell pellets were resuspended in liquid YP medium without supplementation. The OD600 was measured and standardized for each strain such that each well in a Nunc™ MicroWell™ 96-Well Microplates (ThermoFisher Scientific, Waltham, #243656) could be inoculated with a starting OD600 of 0.02. Each strain was inoculated once per thymidine concentration ranging from 0 to 250 µg·mL^-1^ thymidine in increments of 50 µg·mL^-1^. Both strain and thymidine concentration in each well were randomized across the plate. Each 96-well plate was incubated in the Synergy2 plate reader configured with a program maintaining 50°C, medium speed continuous agitation, and measuring endpoint absorbance at 600 nm of each well every 5 minutes for a 24-h period. The growth curve 96-well plate assay was repeated a total of three times on independent days. Growth curve data analyzed as described before.

### Spurious editing analysis

The full base-edited lineages of each EHA105—EHA105, PEN202/EHA105 *thyA*(Q151*), and PEN204/EHA105 *thyA*(Q151*) *recA*(Q178*) and K599 pRi2659ΔT- DNA— K599 pRi2659ΔT-DNA, PEN502/K599 pRi2659ΔT-DNA *thyA*(Q151*), PEN504/K599 pRi2659ΔT-DNA *thyA*(Q151*) *recA*(Q208*) were sent for whole genome sequencing via Plasmidsaurus using Oxford nanopore long-read sequencing.

A *de novo* genome assembly was generated for each the parental EHA105 and K599 pRi2659ΔT-DNA strains using Unicycler v0.5.0 [28]. The overall structure of the *de novo* assemblies were validated using bandage v0.8.1 [29]. The consensus genomes were annotated using PROKKA v1.14.5 [30]. Long reads were filtered with Filtlong v0.2.1 (https://github.com/rrwick/Filtlong) and aligned to the *de novo* assemblies using minimap2 v2.22 [31]. The bam alignments were then used as input along with the annotated assemblies for variant calling with Snippy v4.6.0 (https://github.com/tseemann/snippy) for each the parental samples and their *thyA*- and *recA*- derivatives. Only new variants relative to the parental strain with a read depth >20x were included in the final reported spurious editing frequencies.

### K599 disarming

*A. rhizogenes* strain K599 (NCPPB2659) was disarmed via conventional unmarked allelic replacement using 1kb flanks as described previously for deletion of the *metA* gene in EHA105 and LBA4404 [32].

### Chry5 disarming

*Agrobacterium tumefaciens* strain Chry5 was disarmed via allelic replacement and *flp*- mediated marker excision on a replicating pVS1 binary backbone. The GUS reporter in the FRT-flanked dual *aad*A1/GUS marker cassette from pCPP5243 was exchanged for the sfGFP expression cassette used in pSpCas9d-CDA1-sfGFP-SacB-GGPK/pVP141 via PCR and NEBuilder assembly in previous allelic exchange work (unpublished). The FRT-flanked tandem spectinomycin/sfGFP marker cassette was amplified using overlapping primers to be assembled between 1-kb amplicons from Chry5 that were externally flanking the pTiChry5 T-DNA borders. The three PCR products were isolated as before, and 1 µL of each was combined with 1 µL of *Hind*III and *Eco*RV digested pVPHDB-KAe/pVP075 normalized to 50 ng·µL^-1^ and assembled using 2.5 µL of NEBuilder before transforming into chemically competent *E. coli* 10-beta cells, generating pVPHDBk-efor-C5L-sfGFP/pVP128.

The allelic replacement construct pVPHDBk-efor-C5L-sfGFP/pVP128 was introduced into electrocompetent cells of *A. tumefaciens* Chry5 and plated on YP medium supplemented with 100 µg·mL^-1^ spectinomycin and incubated at 28°C for three days. Eight colonies from the electroporation plate were inoculated into liquid YM (Yeast Mannitol) medium cultures for overnight incubation at 28°C with agitation at 240 rpm in the absence of selection. The next day, 200 µL of the overnight cultures were streaked onto M9 minimal growth medium [33] plates supplemented with 100 µg·mL^-1^ spectinomycin and 10% sucrose and incubated at 28°C to isolate single colonies.

Colonies emerging on the sucrose plate expressing only the visual reporter between deletion flanks (sfGFP), and not the visual reporter on the plasmid backbone (eforRed) were re-streaked on M9 plates supplemented with 100 mg·L^-1^ spectinomycin and 10% sucrose. Colonies on the re-streaked plates were PCR-verified to have the T-DNA region deleted by using primers 5’-GAACATCGGCTGGAATTCGC-3’ and 5’- GTTTCATCAGCCGCGGTTAC-3’ situated outside the deletion flanks. Colony screening was performed with Apex Taq RED Master Mix (Genesee Scientific, El Cajon, CA, #42-138) using the recommended three-step amplification conditions.

For *flp*-mediated excision of the *aad*A1/sfGFP marker from the disarmed pTiChry5, the unstable *flp* expression vector pCPP5264 was introduced by conjugation [34]. Resistant transconjugants growing on LB plates supplemented with 2 µg·mL^-1^ tetracycline were colony-screened using the same primers as used for the initial deletion to confirm ejection of the FRT-flanked cassettes with a smaller 2384 bp, amplicon. Individual colonies were inoculated into liquid LB cultures without antibiotics for overnight incubation at 28°C with agitation at 240 rpm. The following day, 200 µL of the overnight cultures were streaked on LB without antibiotics and incubated at 28°C for three days. Colonies growing on the LB plates were replicate-plated onto LB/LB supplemented with 2 µg·mL^-1^ tetracycline to confirm loss of the *flp* expression vector. The resulting strain possessed a marker-free disarmed pTiChry5 in its native chromosomal background and was named PEN400. The strain was sent for whole genome long-read sequencing via Plasmidsaurus to confirm targets for genome editing and verify the deletion.

To phenotypically support the PCR and NGS deletion, overnight imbibed cotyledons of soybean, *Glycine max*, cv. Jack were separated from the embryonic axis and scored longitudinally on the adaxial surface with a scalpel blade dipped in the mucilage of a 3- day old plate of either wild type Chry5 or disarmed variant PEN400. Cotyledons were then placed adaxial side up in 100 x 15 mm Petri dishes containing an autoclaved Whatman® Grade 1 qualitative filter paper (Cytiva, Marlborough, MA, USA), wetted with 2 mL of liquid ½ MS medium at pH 5.4 and supplemented with 3% sucrose and 100 µg·mL^-1^ acetosyringone (Millipore sigma, Burlington, MA, #D134406). Plates were sealed with Parafilm (ThermoFisher Scientific, Waltham, #1337410), and incubated at 24°C under 16-h light/8-h dark 1-6 µE m^-1^s^-2^ for three days for cocultivation. Then, cotyledons were directly transferred to HRG medium [35] and grown under the same conditions for four weeks before phenotyping. Three biological replicates, blocked by day, were used with 10 cotyledons per treatment.

### Confirmation of virulence of thy- K599

Virulence of the thymidine auxotroph of strain K599 (NCPPB2659) was validated through a modified hairy root assay expanded from that of the Stupar Lab at the University of Minnesota [36]. Briefly, overnight imbibed cotyledons of soybean, cv. Jack were cut parallel to the embryonic axis, and the axis was discarded. Single colonies of wild type K599 and PEN522 (K599 *thyA*(Q151*)) carrying a dicot RUBY expression plasmid (Supplementary Figure 5) were inoculated into 25-mL baffled Erlenmeyer flasks containing liquid YP medium supplemented with 50 µg·mL^-1^ kanamycin and 150 µg·mL^-1^ thymidine and incubated overnight at 28°C with agitation at 240 rpm. The following day, cultures were pelleted in a 4°C refrigerated centrifuge at 4500 x *g* for 30 minutes and resuspended to a final OD600 of 0.5 in liquid ½ MS medium at pH 5.4 supplemented with 3% sucrose and 100 µg×mL^-1^ acetosyringone.

Cut cotyledons were added to sterile polypropylene 50 mL conical tubes, 20 mL of *Agrobacterium* suspension was added and, the conical tubes were placed under vacuum at 80 kPa for 15 min. After 30 total minutes, the *Agrobacterium* suspension was decanted, and the cotyledons were handled as in the Chry5 galling assay. After 4 weeks, cotyledons were scored for expression of RUBY and RUBY hairy roots. The experimental design consisted of three biological replicates blocked by day, with 40 cotyledons per treatment.

## Results and Discussion

### Strain disarming and functional validation

The use of *Agrobacterium spp*. as a genetic engineering tool is the most popular method for transgene delivery into plants. To avoid the disease phenotypes *Agrobacterium* naturally cause, strains are disarmed to eliminate their ability to cause disease and instead deliver genes of interest. Two underutilized strains for crop genetic engineering are K599 and Chry5, each of which have previously demonstrated utility for plant transformation [37, 38]. While K599 has previously been disarmed by other groups [37, 39], we produced our own disarmed variant through allelic replacement to support plant transformation efforts and facilitate subsequent editing of the *thyA* and *recA* loci.

Initially, we attempted to disarm both K599 and Chry5 using conventional allelic replacement in a similar manner to how Prias-Blanco et al., (2022) generated methionine auxotrophs of EHA105 and LBA4404 [32]. We used this method to successfully disarm K599, as confirmed via PCR and NGS sequencing. However, disarming pTiChry5 was more difficult, and a line with the desired allelic replacement could not be recovered after screening hundreds of isolates. Therefore, we adopted a modified marker excision strategy expanded from [34]. Our modified strategy sought to incorporate two unique antibiotic resistances and two unique chromoproteins in tandem to allow for efficient visual and chemical screening of desirable recombination events.

The vector pCPP5243 [34] encodes an FRT-flanked dual *aadA1*/GUS marker cassette for marked selection. The dual marker can later be ejected through introduction of a plasmid expressing the *flp* recombinase. The GUS reporter, however, requires X-Gluc to be provided in the medium for visual selection. We opted to exchange the GUS reporter for sfGFP [40] to simplify the deletion workflow and medium components.

Furthermore, as initial attempts at using a non-replicative origin on the allelic replacement/disarming plasmid failed to produce any single or double crossover events in Chry5, and since we were using markers between deletion flanks, we cloned our deletion cassette onto the pVS1 vector backbone pVPHDB-Se/pVP073 capable of replication in *Agrobacterium* which was itself marked with a separate antibiotic resistance gene and chromoprotein. While using a replicating plasmid for allelic replacement can result in increased persistence of colonies without crossover events, it was the method originally used in the construction of Chry5 derivative, KYRT1 [38]. By using two sets of chromoprotein-antibiotic resistance genes delivered in tandem, one between deletion flanks and the other on the vector backbone, we were able to efficiently screen double crossover events via the loss of the backbone markers. Once cell lines that had lost the backbone markers were isolated, we introduced an unstable *flp* expression plasmid [34] to excise the dual marker that had been recombined between the flanks. Marker excision was scored for loss of sfGFP expression, screened via PCR (Figure 2B), and confirmed via long-read Oxford Nanopore NGS sequencing. To further validate the deletion through phenotyping, we infected overnight imbibed cotyledons of soybean, *Glycine max* cv. ‘Jack’, and scored them with each wild type Chry5 and our disarmed variant PEN400. Four weeks after infection, cotyledons were phenotyped (Figure 2C). All cotyledons infected with the armed strain expressed varying degrees of galling, while no cotyledons infected with the disarmed strain galled, further validating that our allelic replacement strategy was effective.

**Figure 2.**
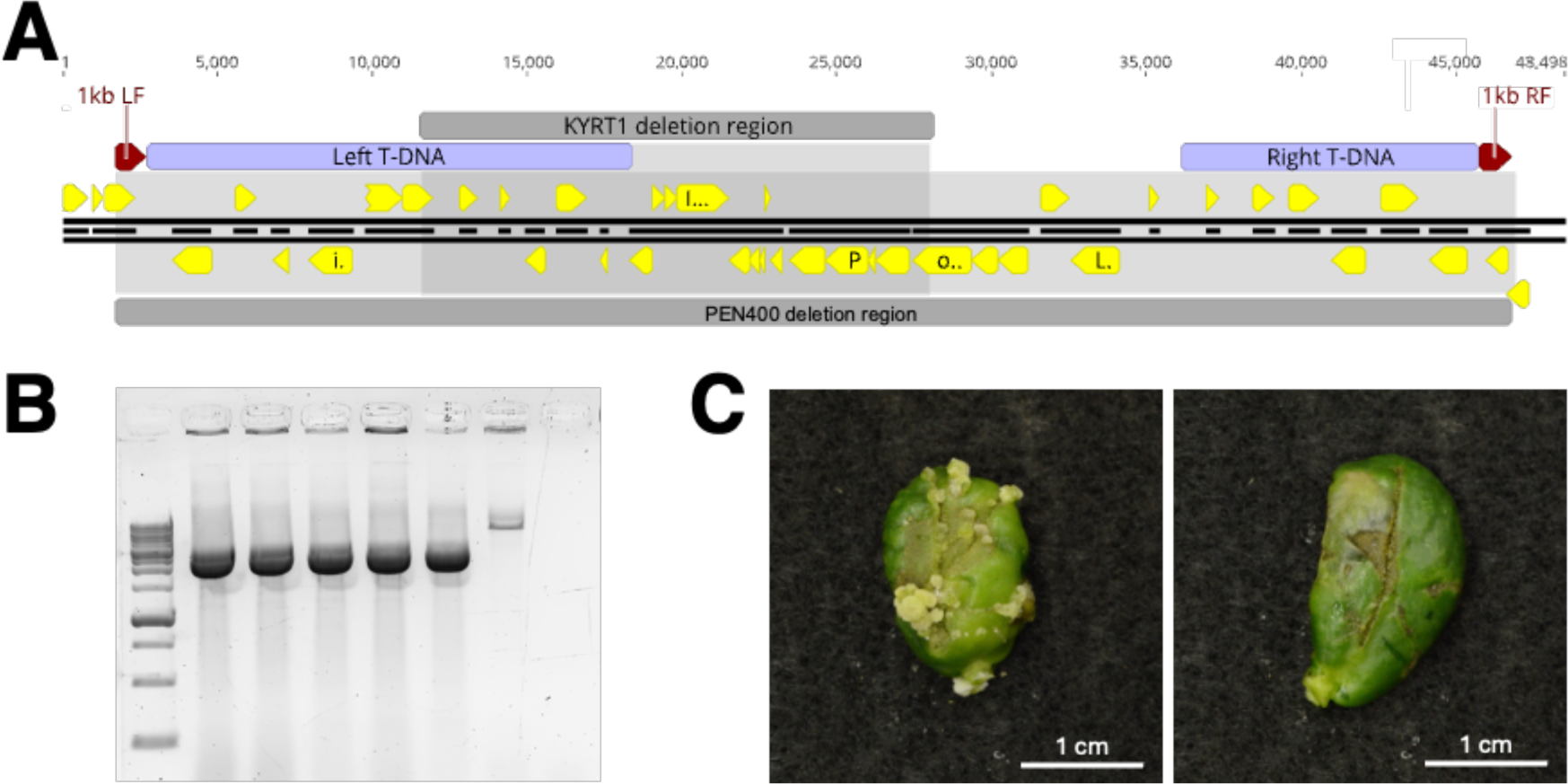
Disarming *Agrobacterium tumefaciens* Chry5. A) T-DNA region of pTiChry5 with annotations for the left T-DNA, right T-DNA, region deleted to generate partially disarmed strain KYRT1[38], left and right deletion flanks to generate PEN400, and the deletion region of PEN400. Yellow annotations represent coding sequences within the displayed portion of pTiChry5. B) Gel electrophoresis of PCR amplicons spanning the marker-ejected deletion region in five independent colonies. L to R: 1 KB ladder, 5 independent colonies of PEN400, positive control of the marked deletion, negative control. C) Soybean cotyledons 4 weeks after infection with wild type Chry5 (left) and disarmed Chry5 (PEN400)(right) confirming loss of its galling ability.

Our fully disarmed and unmarked Chry5 derivative, PEN400, retains a single FRT site on the pTiChry5ΔT-DNA due to the *flp*-mediated marker excision. This FRT site may prove useful should co-integrating vectors be desired to express T-DNAs containing genes of interest from the native Ti plasmid instead of in a binary system. We expanded our disarmed Chry5 lineage further by introducing each thymidine auxotrophy and thymidine auxotrophy stacked with *recA* deficiency (Table 1) to better serve the plant transformation community.

### Agrobacterium mutant generation with single component base editors

Previously established CRISPR-mediated base editors for gene knockout in phytopathogenic bacteria have been the products of successive restriction ligation and Gateway assembly of four modules [19, 20]. The modules correspond to a plasmid backbone capable of replication in the host organism, a *sacB* expression cassette for counterselection upon plating on sucrose to evict the editing plasmid, a gRNA expression cassette, and the editor expression cassette. Given that these components function together reliably, we elected to combine them into one plasmid that guides can be cloned into, obviating the need for a Gateway assembly step.

We initially reconstituted the final base editor vector for engineering *Agrobacterium* 1. [19] as a single component plasmid. Rather than restriction-ligate a gRNA into a component module, and later recombine that module using Gateway recombination into a final editor plasmid, the gRNA cloning takes place directly in the final vector. Our initial editor plasmid, pSpCas9d-CDA1-sfGFP-SacB-GGPK/pVP141, essentially reconstituted the final Gateway product of [19], as if a gRNA had never been cloned into its Gateway entry module. It also exchanges spectinomycin resistance for kanamycin resistance.

This editor plasmid is a useful tool to recapitulate the system already available in the literature, but isolation of initial edited lines highlighted another challenge to using base-editor vectors: plasmid eviction. Plasmid eviction on all base editor plasmids for phytopathogenic bacteria to date employ the *sacB* marker for counterselection on sucrose, and counterselection is an important feature to ensure that the editor is only present for a finite duration to make the desired edit. Our initial cell line isolated after targeting the thymidylate synthase, *thyA*, in strain K599 pRi2659ΔT-DNA failed to evict the editor plasmid when plated on sucrose. This error was not immediately apparent. We therefore sought to produce two new editor plasmids that introduce a chromoprotein into the vector backbone, enabling visual monitoring of plasmid eviction under ambient conditions without any added expense or molecular screening burden. Single component base-editors pSpCas9d-CDA1-sfGFP-SacB-KAe/pVP143 and pSpCas9d-CDA1-sfGFP-SacB-Se/pVP144 were generated, expressing each the AeBlue chromoprotein [27] and kanamycin resistance in pVP143, and the eforRed chromoprotein [27] and spectinomycin resistance in pVP144, respectively. These two additional single component base editors simplify the mutant generation process by 1) eliminating Gateway recombination for assembly, and 2) enabling visual monitoring of editor eviction on sucrose without requiring additional consumables or UV lighting.

We next compared the eviction frequency for each chromoprotein-marked editor backbone by comparing both chromoprotein marked editors in EHA105, LBA4404, and K599. Only the pVP144 derived editor was tested in strain Chry5, as it has a high level of basal kanamycin resistance that precludes the use of pVP143-derived editors. Eviction frequency was high for the editors in all backgrounds with an overall average of 92 ± 8%. However, escapes were also present in all strain backgrounds, highlighting the importance of being able to easily screen for loss of the editor plasmid.

To further simplify the use of base-editors for generating mutant strains of *Agrobacterium*, we developed a simple Python script named the Base Editor Enrichment Function, or BEEF, for filtering a list of candidate gRNAs, and exporting only those that can produce a premature stop codon. BEEF is executable from the command line on either MacOS or Windows. We also connected the script to a Geneious Prime wrapper plugin, named PrimeGradeBeef. To execute the script through Geneious Prime, users can install the PrimeGradeBeef plugin and associated Geneious Prime workflow, named The Butcher. The Butcher executes the underlying Python script on a Geneious file containing only an annotated coding sequence (Supplementary Figure 2A). The workflow will prompt the user to select a locally stored database for off-target scoring (B), and then execute the Doench et al. (2016) guide prediction algorithm for 20-nt guides. The Butcher will automatically pass the output of Doench et al. (2016) to the underlying Python script, filter the candidate gRNAs, and output a file containing only the coding sequence and associated viable guide annotations (Supplementary Figure 2C). The end user can then select specific guides for gene knockout based on each guide’s location within the coding sequence, as well as the predicted off target binding. The plugin and workflow are designed specifically for the Target-AID architecture but can be modified to accommodate other architectures.

Alternatively, wrapper plugins for guide filtering with alternative software can be designed.

As proof of concept, and to expand the available mutants of laboratory *Agrobacterium* strains in the public domain, we used our simplified workflow and editor plasmids for CRISPR-mediated base-editing to introduce thymidine auxotrophy and *recA* deficiency in several strain backgrounds (Table 1). Each mutation was chosen for its specific benefit posed to plant transformation—easier elimination after infection and maintained plasmid fidelity in *Agrobacterium* [12, 41]. An example base edit of a thymidylate synthase target can be seen in Figure 4A. Thymidine auxotrophy leads to near-complete inhibition of bacterial growth, even in nutrient-rich media such as LB, provided no exogenous thymidine is supplied (Figure 4B). Since the *recA* mutations were produced serially after *thyA* mutation, we whole genome sequenced our lineages of EHA105 and K599 pRi2659ΔT-DNA derivatives to evaluate spurious editing. Spurious editing was observed in each base-edited sample at a minimum read depth of 20x, with increased spurious editing observed in the *recA* minus mutant at that same cutoff (Supplementary Figure 3).

**Figure 3.**
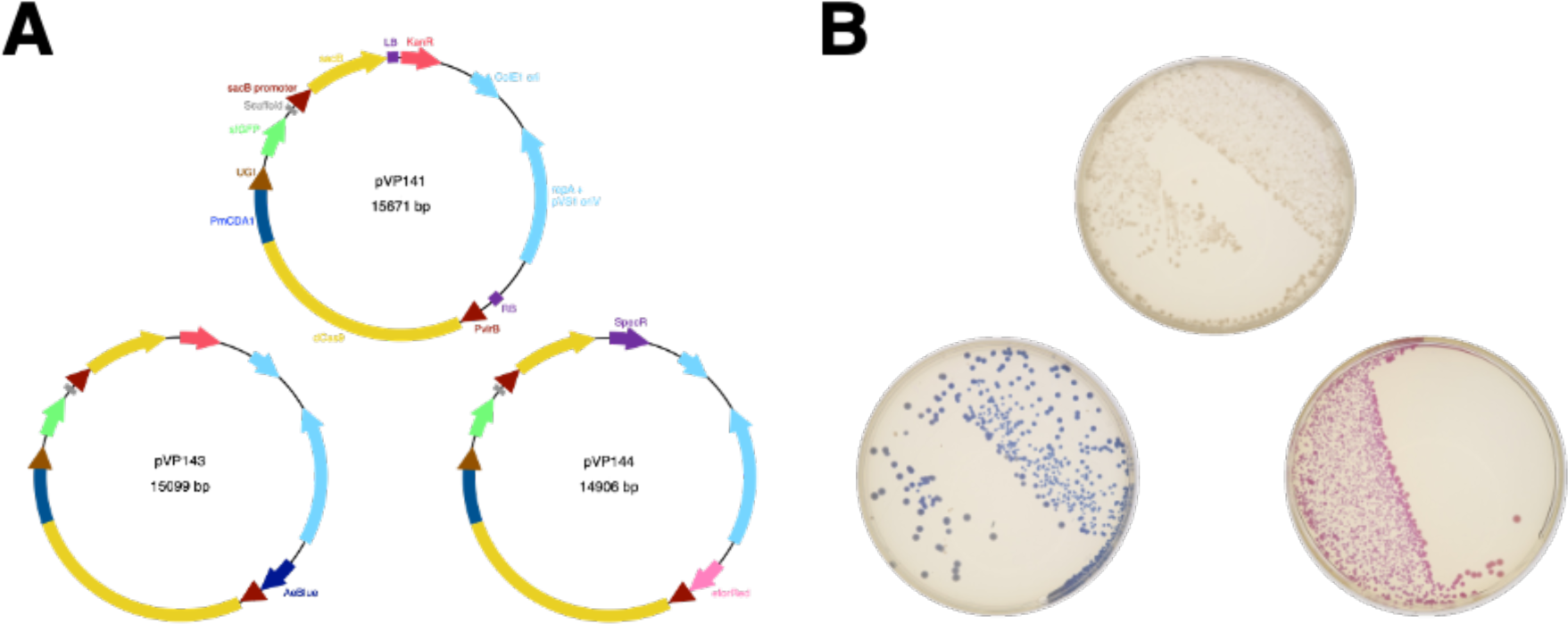
Single component base editors for generating base-edited *Agrobacteria*. A) Three single component base editor plasmids were produced. The first, pVP141 is a kanamycin-resistant reconstitution of the Gateway product from Rodrigues, Karimi [19]. This vector contains sfGFP for visualization of marker excision when cloning sgRNAs into the plasmid. The second single component editor plasmid, pVP143, removes the unnecessary T-DNA borders form the original plasmid, and adds a second chromoprotein, AeBlue, to the backbone. The AeBlue chromoprotein in the backbone enables visual selection against colonies that fail to evict the editor plasmid when plated on sucrose. Finally, pVP144 exchanges kanamycin resistance for spectinomycin resistance, and the AeBlue chromoprotein for eforRed, enabling use in kanamycin-resistant backgrounds. B) Representative plates of what each editor plasmid looks like under ambient light in *A. rhizogenes* strain K599 7 days after electroporation.

**Figure 4.**
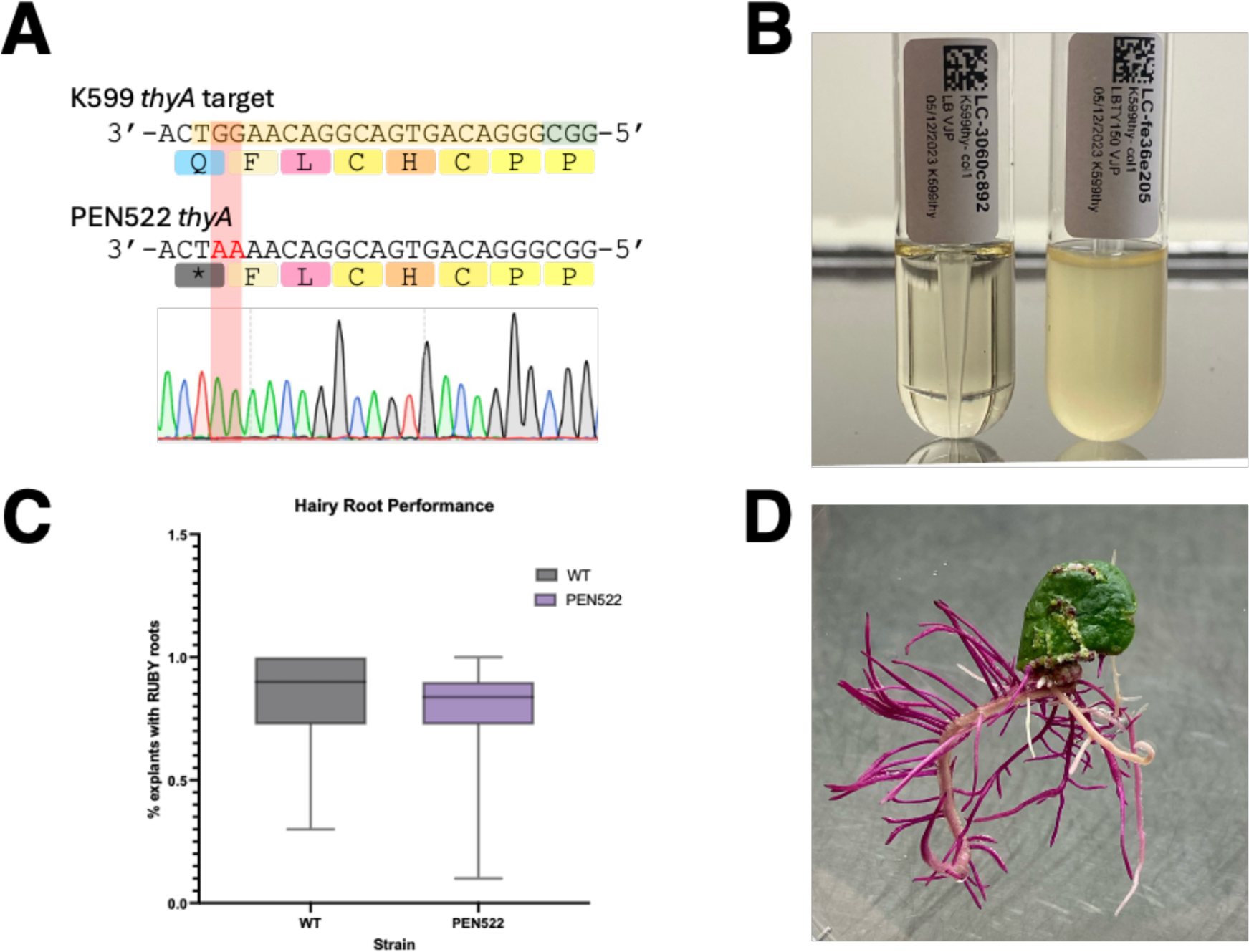
Example editing in *Agrobacterium rhizogenes* strain K599 (NCPPB2659). A) Chromatogram displaying the knockout of the thymidylate synthase gene in K599 (NCPPB2659). The cell line recovered possessed the desired Q151* mutation, effectively knocking out function of thymidylate synthase. B) Thymidine-dependent growth of strain PEN522 (K599 *thyA*- (Q151*)) after 4 days of incubation at 28°C with shaking. C) Hairy root generation comparison of thymidine auxotrophic (PEN522) vs wild type (K599) carrying a dicot RUBY reporter plasmid. D) Representative example of hairy roots expressing RUBY 4 weeks post infection.

To ensure there was no loss of virulence after initial base editing, strain PEN522 (K599 *thyA*(Q151*)) was compared against its wild-type counterpart K599 carrying a RUBY reporter plasmid in a replicated hairy rooting assay (Figure 4C). Data were subjected to an unpaired t-test, demonstrating no significant difference between strain performance under the experimental conditions tested (p=0.81), suggesting the base-edit had negligible impact on transformation efficiency, measured as the proportion of explants expressing RUBY hairy roots.

### Agrobacterium strain performance

We evaluated if there were fitness costs associated with thymidine auxotrophy and *recA* deficiency in terms of bacterial growth as previously suggested [42, 43]. When our suite of edited strains was isolated, fitness costs were only noted in some *recA-* backgrounds, most notably K599 (Figure 5B, Figure 6B). To explore fitness costs further, we assayed our EHA105, K599 pRi2659ΔT-DNA, and PEN400 edited lineages, which represent three distinct chromosomal backgrounds (Figure 5). LBA4404 was not assayed as its aggregating behavior in liquid culture interferes with spectrophotometric measurements. The only response difference for thymidine auxotrophy can be seen in the top panel of Figure 5B where there appears to be a growth lag and reduction in OD during extended culture of PEN502 (ΔK599 *thyA-* (Q151*)) relative to its wildtype counterpart at all thymidine concentrations, but particularly at 50 mg·L^-1^ thymidine.

**Figure 5.**
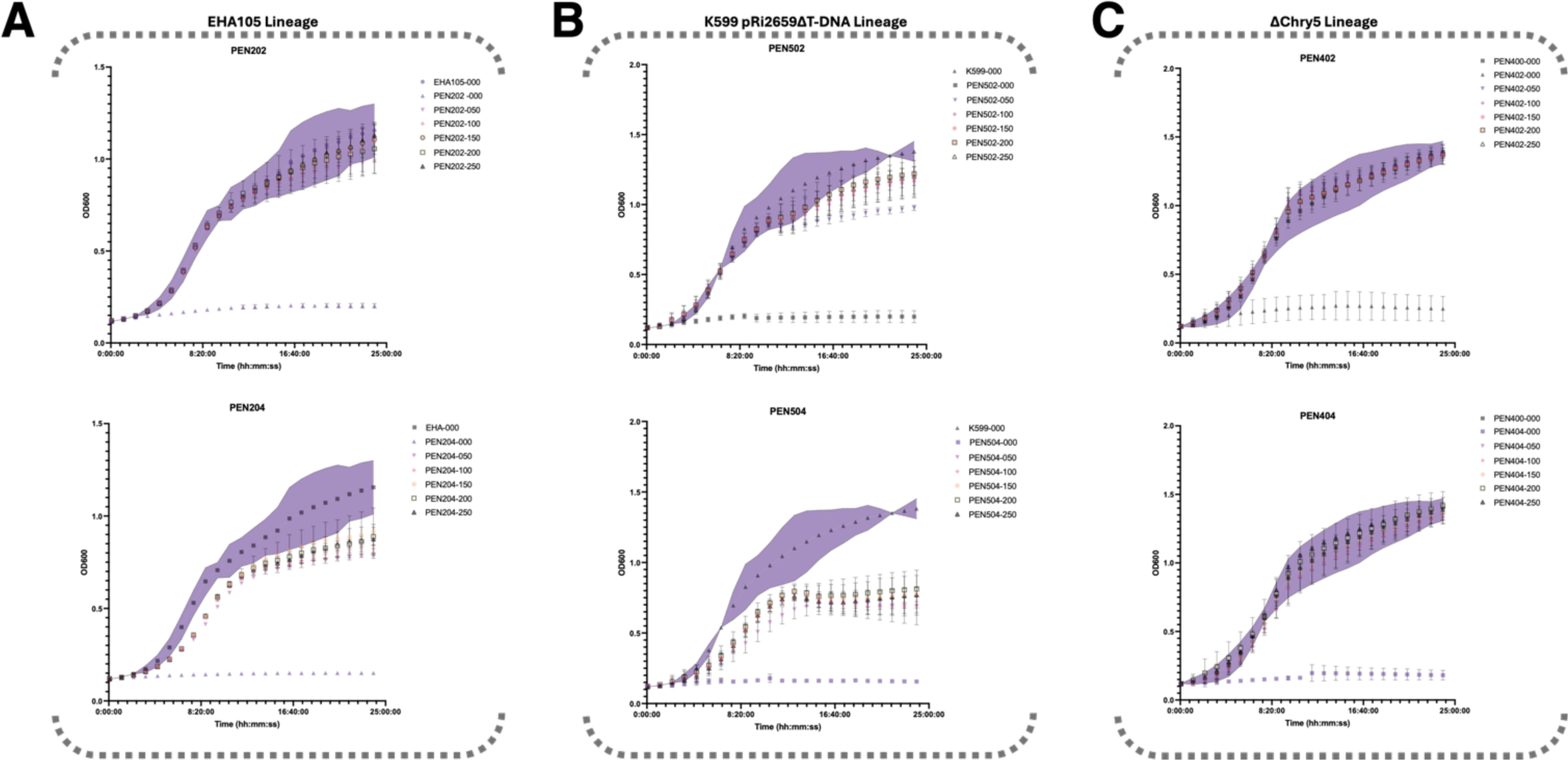
Growth curves corresponding to the A) EHA105, B) K599 pRi2659ΔT-DNA and C) PEN400 strains. Purple 95% confidence intervals are displayed on each chart for each wild-type strain. The data represent three biological replicates blocked by day and grown in 96-well plate format measuring OD600 every 5 minutes for 24 hours. Staring OD was normalized to 0.02 per well. Legend keys use strain names hyphenated with a number code in the plot legends indicates the concentration of supplemented thymidine in the medium in mg×L^-1^ thymidine.

**Figure 6.**
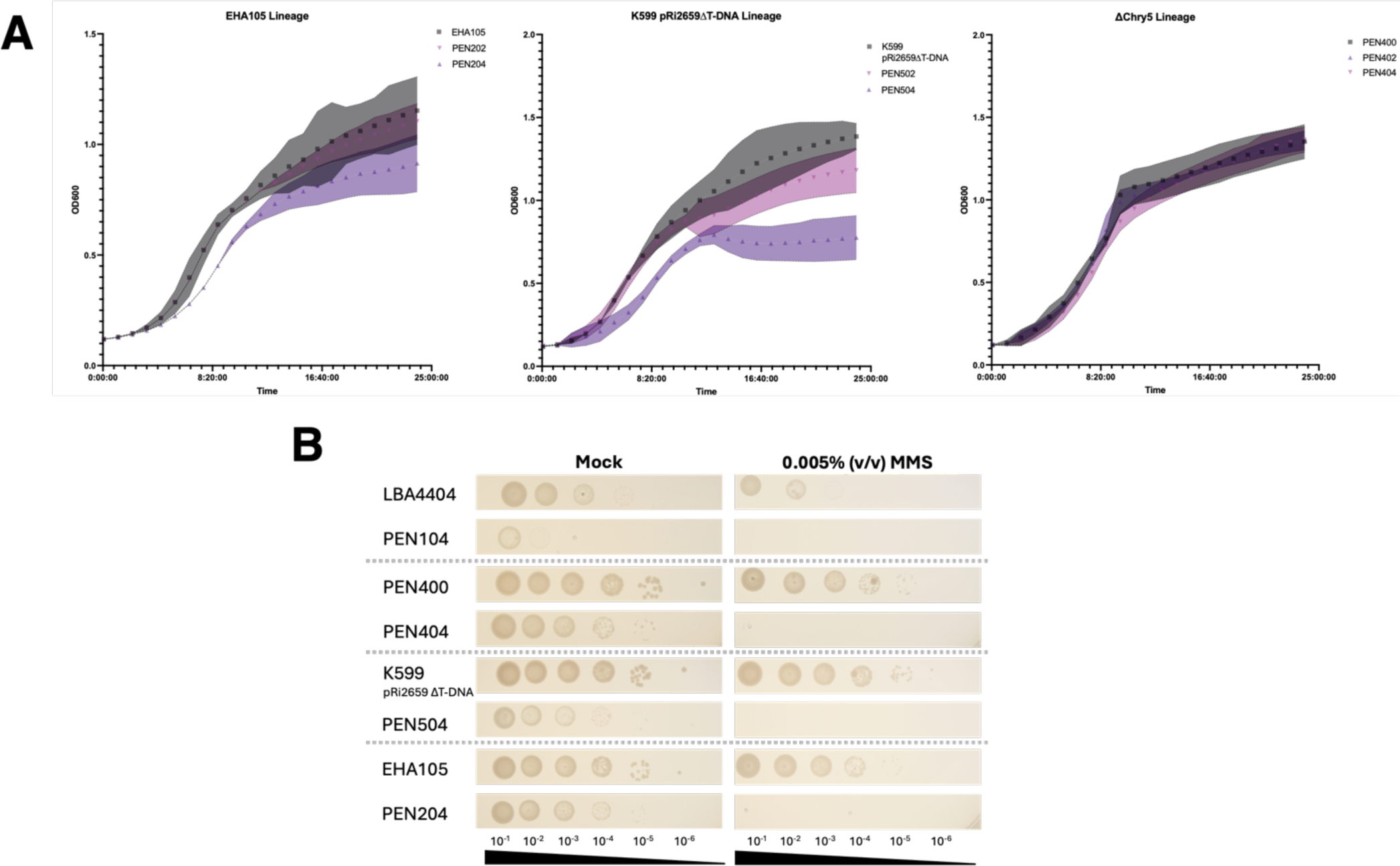
Evaluation of growth of *recA- Agrobacterium* mutants. A) Growth curves for edited EHA105, K599 pRi2659ΔT-DNA, and PEN400, highlighting differences in growth in liquid YP medium supplemented with 150 mg×L^-1^ thymidine. Growth curves were collected as before, and confidence intervals represent the standard deviation of three biological replicates. B) Six serial dilutions of overnight liquid cultures for each LBA4404 *recA ±*, Chry5 pTiChry5ΔT-DNA *recA ±*, K599 pRi2659ΔT-DNA *recA ±*, and EHA105 *recA ±*. Serial dilutions were replicate-plated on solid YP medium (negative control) and YP medium supplemented with 0.005% (v/v) MMS and 150 mg×L^-1^ thymidine to evaluate DNA-damage sensitivity conferred by knockout of *recA*. Plates were imaged after incubation for 48h at 28°C.

The *recA* mutation, however, causes a fitness cost in the K599 pRi2659ΔT-DNA background, accounting for nearly a 43% reduction in OD during stationary phase. Meanwhile, it results in only a 24% reduction in stationary phase OD in the EHA105 background, and no detectable fitness cost in the PEN400 background (Figure 5). The lack of a fitness cost of *recA* mutation in the PEN400 background was unexpected and requires further exploration (Figure 6).

To further evaluate the potential fitness costs, or lack thereof, of the *recA* minus trait in the ΔChry5 background, we plotted growth curves for the EHA105, K599 pRi2659ΔT- DNA, and PEN400 stains at a fixed thymidine concentration (Figure 6A), and obtained similar results.

To assess whether the *recA* minus strains are truly devoid of *recA* activity, we performed a methyl methanesulfonate, or MMS, screen on each of our LBA4404, PEN400, K599 pRi2659ΔT-DNA, and EHA105 lineages (Figure 6B, Supplementary Figure 4). In all lineages, there was complete mortality for strains rendered r*ecA* minus via base-editing that were plated on MMS, while *recA* positive strains exhibited only partially inhibited growth (Figure 6; Supplementary Figure 4). These data, in conjunction with Sanger sequencing, confirmed that the *recA* gene in PEN404 (ΔChry5 *thyA-* (Q151*) *recA*- (Q178*)) has been rendered non-functional through CRISPR-mediated base-editing, although the mutation does not appear to impede growth as it does in other strain backgrounds. The genetic basis for this difference in behavior is not immediately apparent but highlights the importance of having a diverse set of *Agrobacterium* strains available for plant transformation purposes, as the different backgrounds may have unique performance features.

## Conclusions

Previous methods of mutant generation in *Agrobacteirum* were either time consuming, non-specific, or a combination of the two. Allelic replacement in the absence of accessible sequencing, like when the originally disarmed strains were developed, resulted in deletions that were not nearly as specific as can be achieved today.

Previously, auxotrophy, and other mutations that can be readily phenotyped, were isolated after widespread genome disruption through nonspecific mutagenesis. Given recent advancements in gene editing, we can be much more targeted with our approach to generating mutant strains and do so now more quickly than ever before.

We constructed and deployed a series of single-component base-editor vectors and generated a suite of *Agrobacterium* strains with enhanced functionality for laboratory use. We introduced thymidine auxotrophy and *recA* deficiency in the commonly used strains LBA4404, EHA105, and GV3101::pMP90, and developed disarmed versions of *A. rhizogenes* strain K599 and *A. tumefaciens* strain Chry5. Our newly disarmed strains were also edited for both thymidine auxotrophy and *recA* deficiency, further expanding the available lineages of disarmed *Agrobacterium* strains for laboratory use.

To facilitate and support the editing process, we developed a simple Python script and associated Geneious Prime workflow to streamline guide filtering for the Target-AID architecture, simplifying and accelerating the design of new strains via base-editing. Our single component editor plasmids will be made available through Addgene for widespread distribution and the code for the guide filtering Python script is available on GitHub (https://github.com/vincentpennetti/PrimeGradeBeef). Our suite of disarmed and edited strains will be made available for distribution.

## Acknowledgments

We thank Dr. Thomas Jacobs (VIB–Ghent University, Ghent, Belgium) for providing the pEN-L4-PvirB-dCas9-CDA-UL-T3T-R1, pEN-L1-PJ23119-Bsa1-PglpT-sfGFP-TrrfB-Bsa1-Scaf-L2, and pEN-R2-SacB-L3 vectors; Sebastian Cocioba for the promoter-rbs-spacer combination driving eforRed; and Dr. Timothy Chappell for disarming *Agrobacterium rhizogenes* strain K599 through allelic replacement.

## Funding

Funding was provided by the U.S. Department of Energy, Office of Science, Office of Biological and Environmental Research program under Award number DE-SC0023338 and DE-AC05-00OR22725 through the Center for Bioenergy Innovation.

## Competing interests

The authors declare that there is no conflict of interest regarding the publication of this article.

## Data Availability

Raw nanopore sequencing data is available through the NCBI Sequence Read Archive BioProject number PRJNA1143588. Single component editor plasmids and their progenitor backbones will be made available via Addgene (Waltham, MA).

---

**Supplementary Figure 1.**
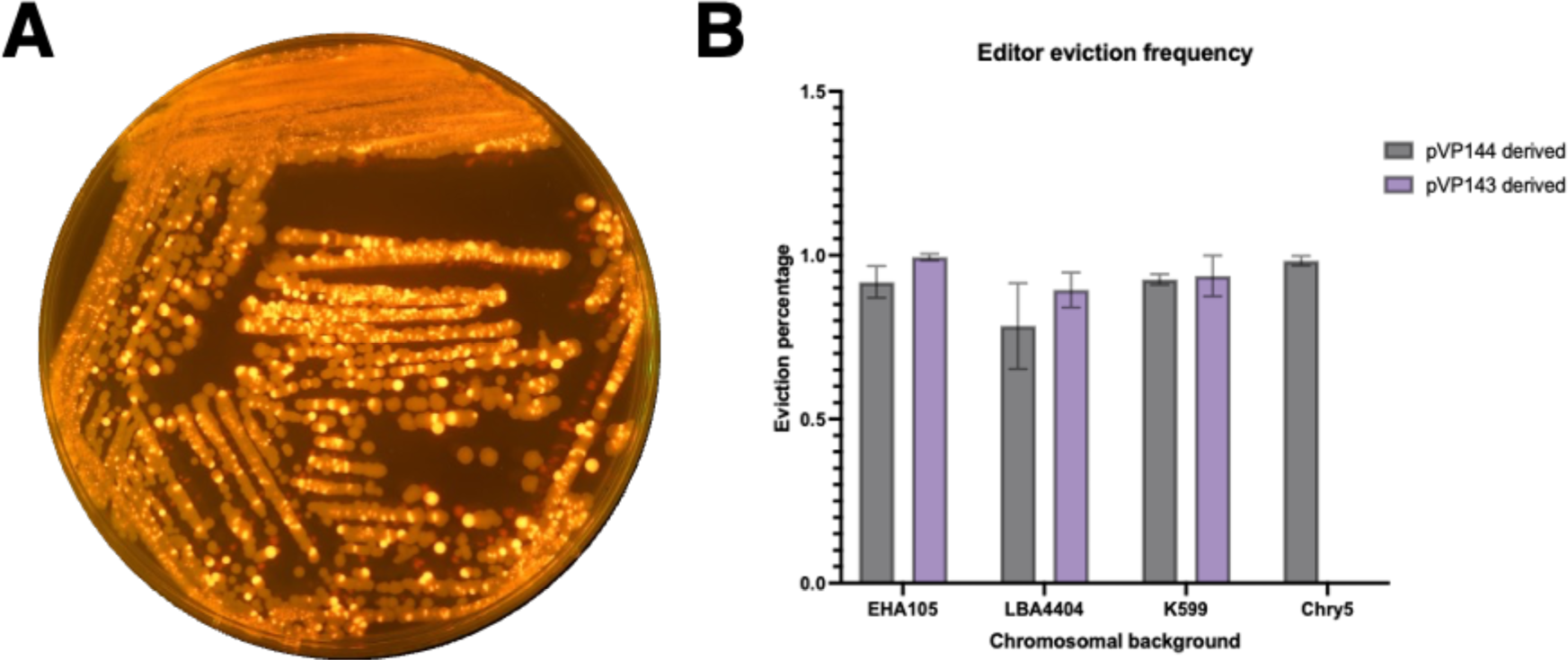
Base editor eviction efficiency. A) A representative colony of a poorly evicting colony of Chry5 containing a pVP144-derived editor targeting the thymidylate synthase gene. Note that the colony was directly streaked from an electroporation plate onto sucrose, without resuspension or overnight incubation, resulting in a particularly poor frequency of eviction. Light orange fluorescing colonies represent colonies that failed to evict the editor plasmid. The colonies also appear pink under ambient light. Dark red spots are glare introduced during transillumination of the Petri dish. B) Eviction data for each strain background based on three biological replicates. Each chromoprotein-marked editor plasmid, pVP143 and pVP144, was tested in each strain background, except Chry5, as Chry5 is incompatible with pVP143 given its high basal kanamycin resistance. When resuspending colonies and incubating overnight prior to eviction, the eviction frequency for each strain is high, but remains imperfect. Mutant generation can be expedited with the chromoprotein-marked backbones by directly plating on sucrose and visually selecting for the colonies that have lost the chromoprotein reporter.

**Supplementary Figure 2.**
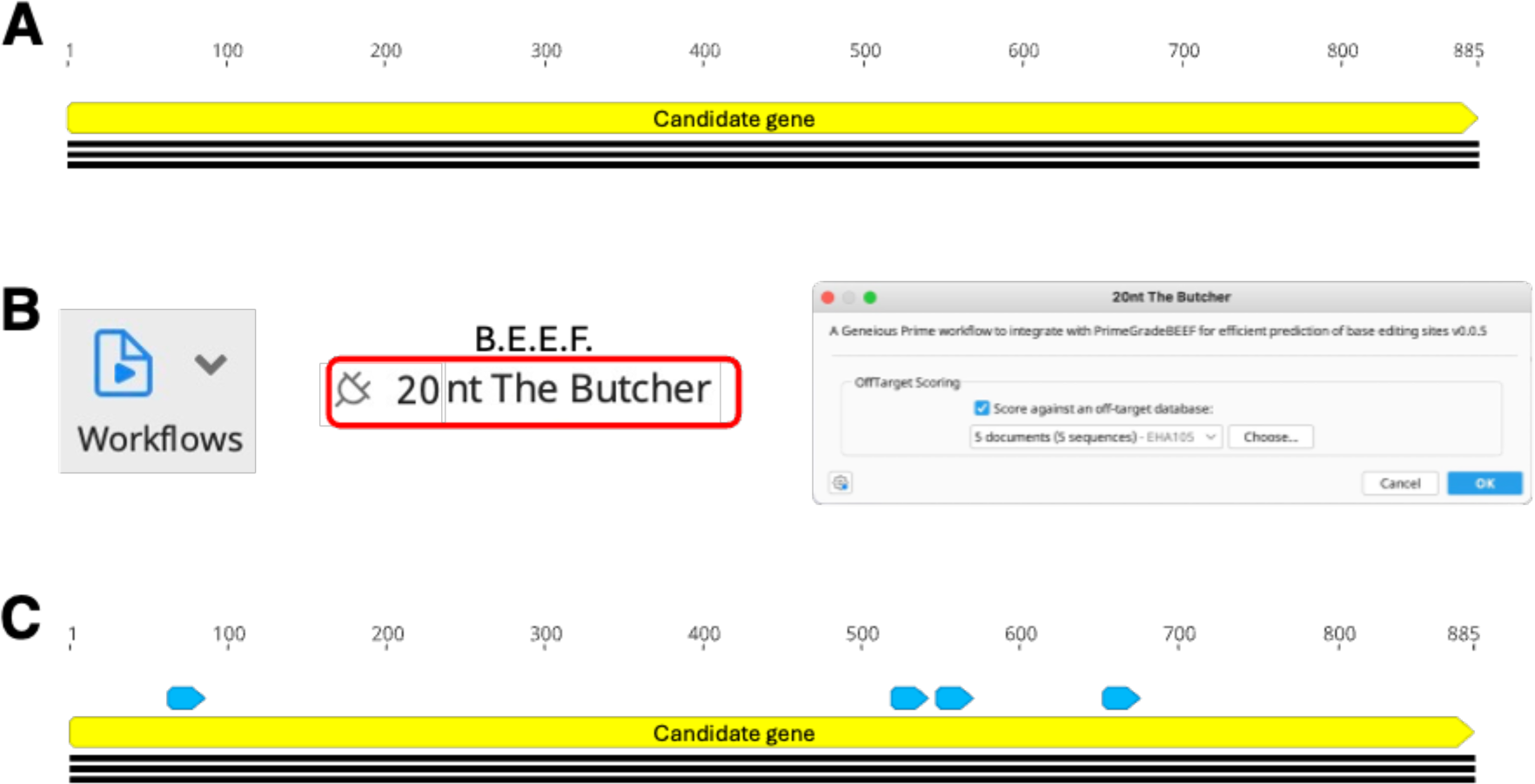
A sample workflow of using PrimeGradeBEEF for guide filtering in Geneious Prime. A) A Geneious file containing the CDS to be knocked out is used as input for automated guide filtering with BEEF. B) Once the CDS file is generated and opened, the workflow ‘20 nt The Butcher’ can be executed for automated guide filtering. The user will be prompted to score against a local off-target database. If this is selected, a subsequent popup (not pictured) will appear, confirming that the candidate gene is already present in the off-target database and will be ignored. C) The Butcher returns a new result file to the user, with putative gRNAs annotated on the map. Each gRNA contains a cytidine residue within the Target-AID editor’s window that can be converted to a thymidine to generate a premature stop codon. Each guide annotation retains the associated off target metadata. Final candidate gRNA for the knockout is user-determined based on location within the CDS and predicted off target binding.

**Supplementary Figure 3.**
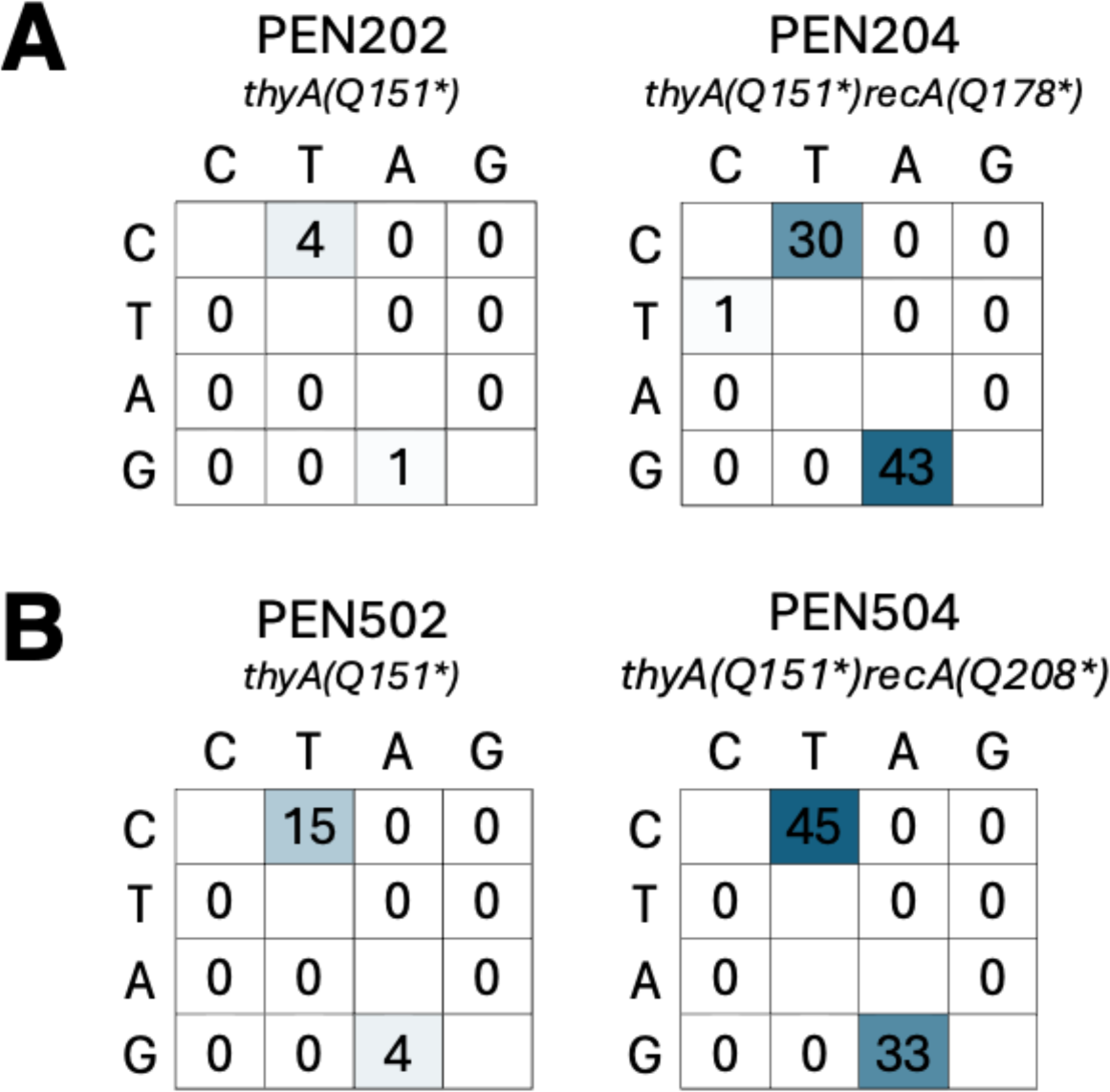
Spurious editing in the EHA105 lineage (A) and K599 lineage (B). Variant calling was performed by mapping long read ONT data against the *de novo* assembled genome sequences. Snippy version 4.6.0 was used for variant calling. Box shading is dependent on value within. Spurious edits are reported as the number of new spurious edits in the sample compared to the progenitor strain. Aligned read depth was cutoff at a minimum of 20x for spurious edit variant calls.

**Supplementary Figure 4.**
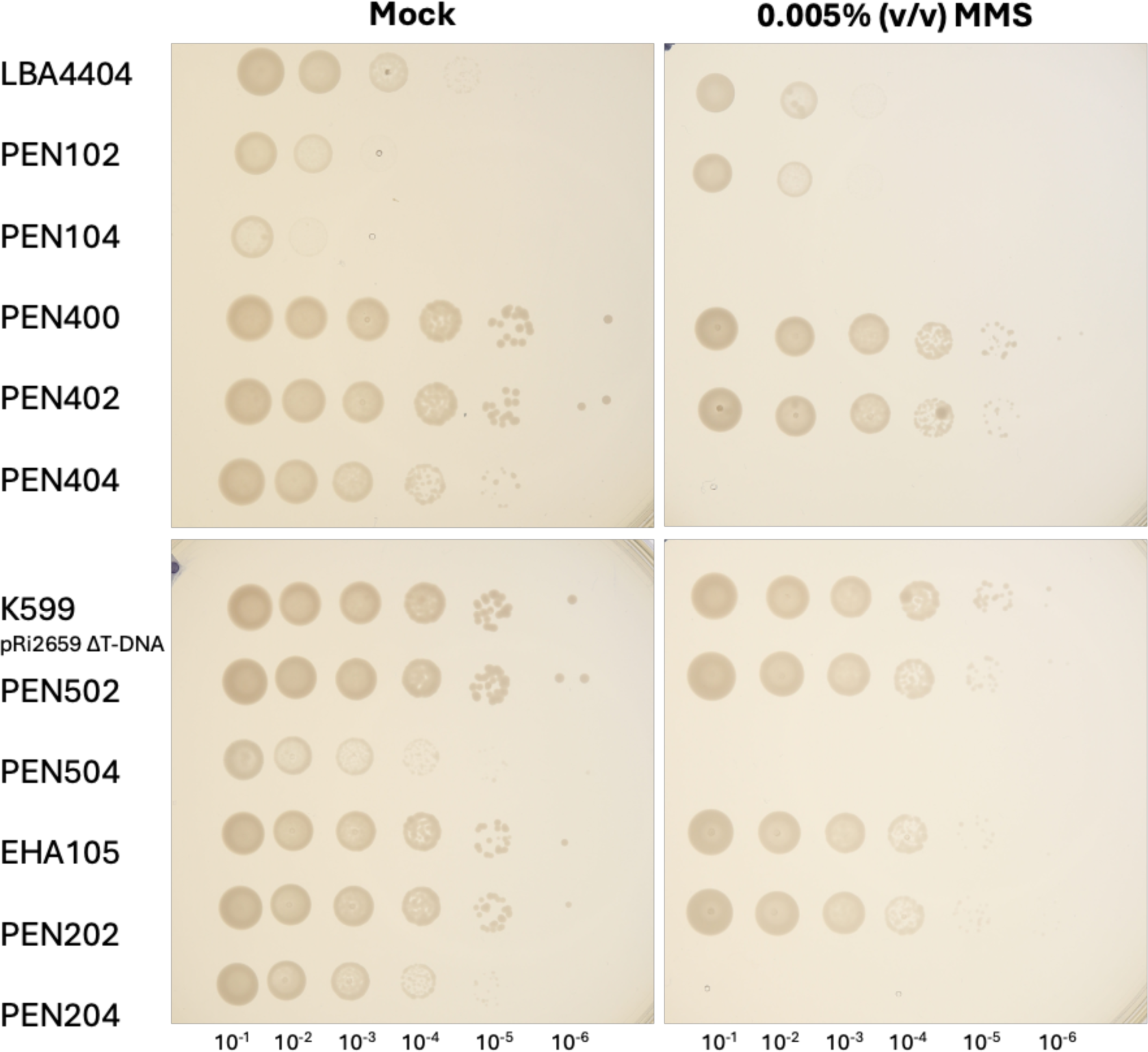
Uncropped image of replica plated MMS screen for primary strain lineages. Six serial dilutions of each strain were plated, representing the full mutant lineage for each LBA4044, PEN400, K599 pRi2659 ΔT-DNA, and EHA105. Plates were imaged after incubation for 48h at 28°C.

**Supplementary Figure 5.**
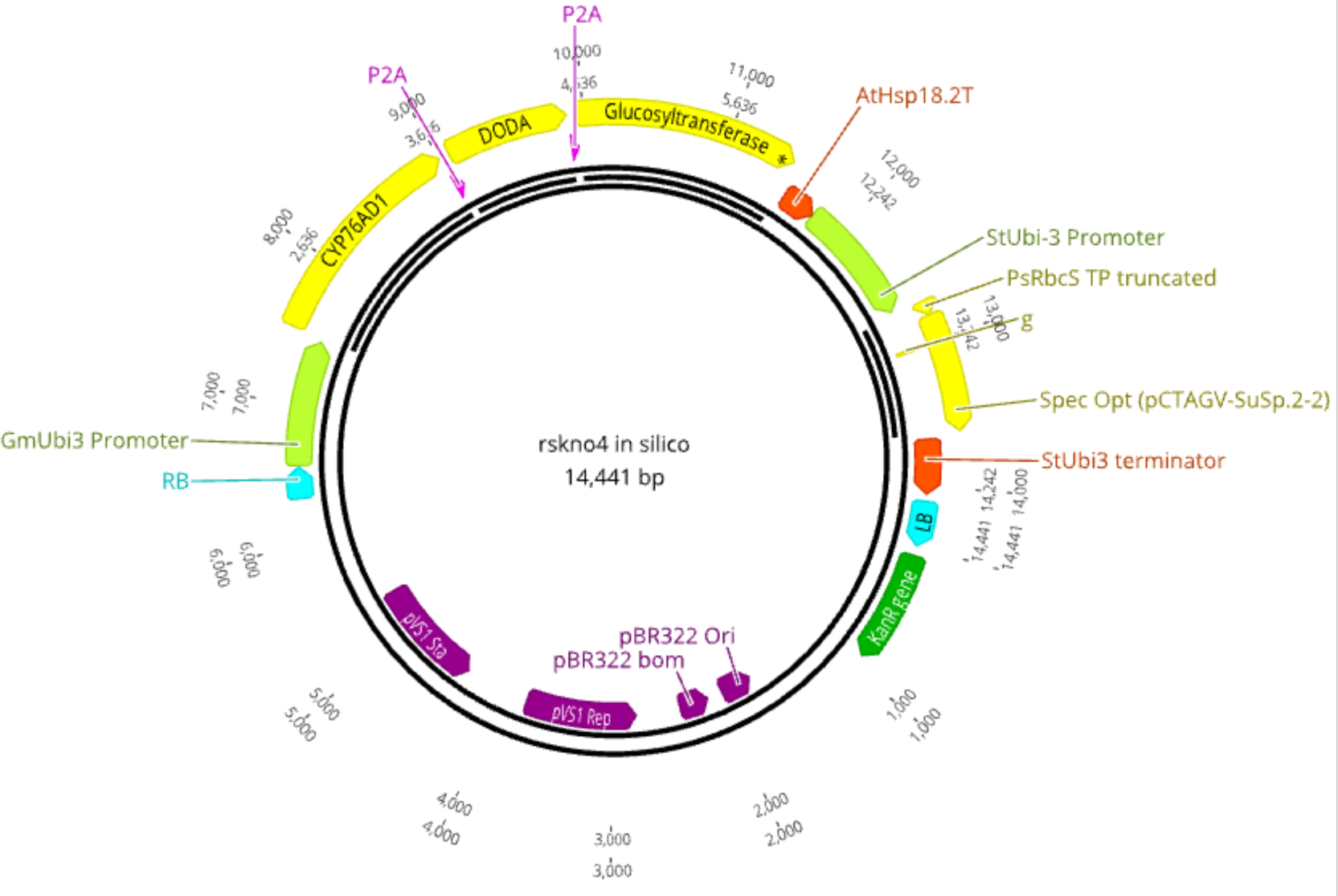
Plasmid map of dicot ruby expression vector. The RUBY visual reporter [45] is driven by each the ubiquitin-3 promoter from *Glycine max* [46] and the heat shock protein 18.2 (AT5G59720.1) terminator from *Arabidopsis thaliana.* Spectinomycin plant selection is conferred from expression of a spectinomycin *aadA*1 resistance gene (Genbank QID24729.1) driven by the ubiquitin 3 promoter and terminator from *Solanum tuberosum* [47]. The assembly is the GreenGate product assembled into vector backbone is pGGPK-AG2 (Pennetti et al., 2024).

**Supplementary Table 1.**
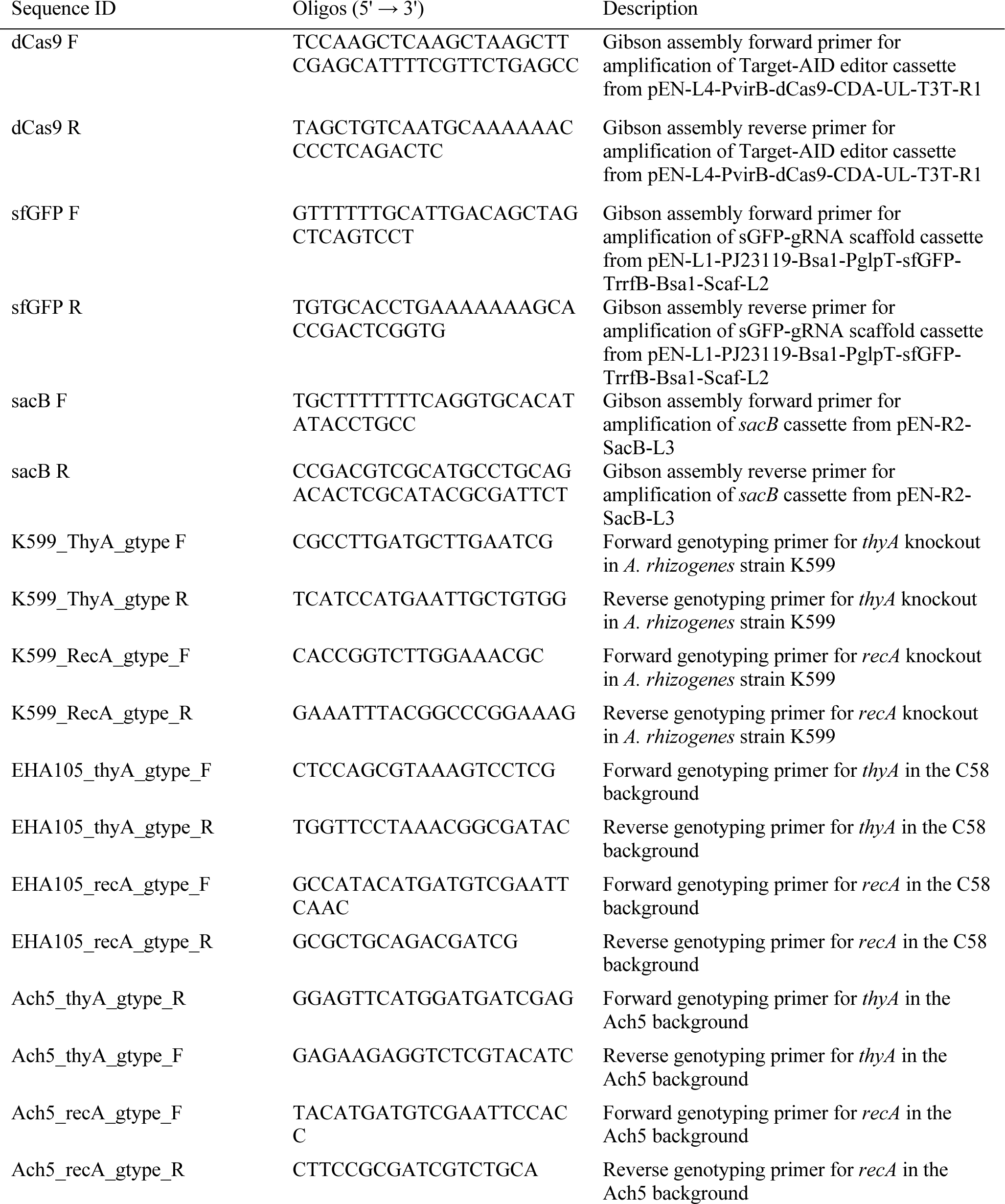
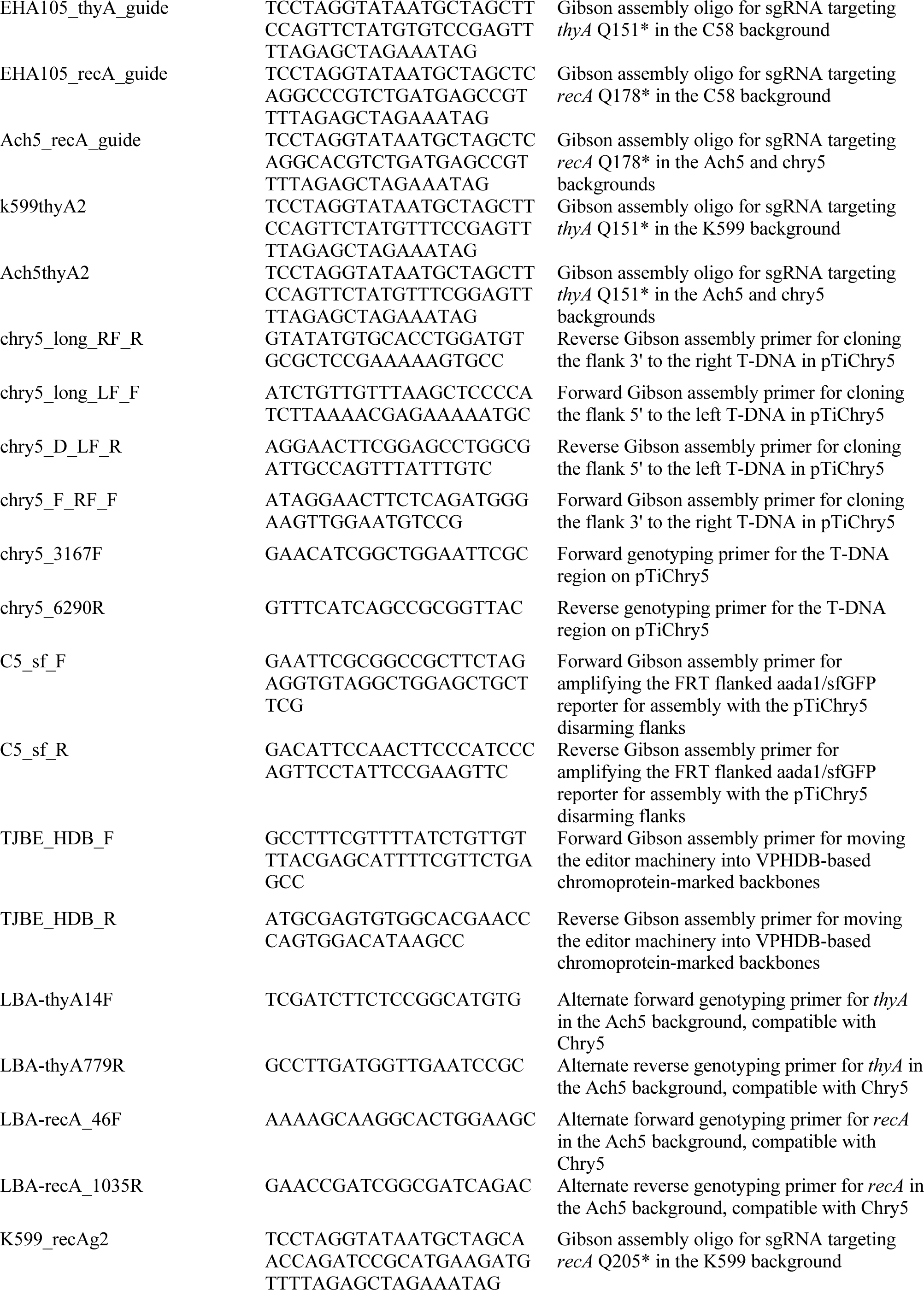
List of primers used in this study.

**Supplementary Table 2.**
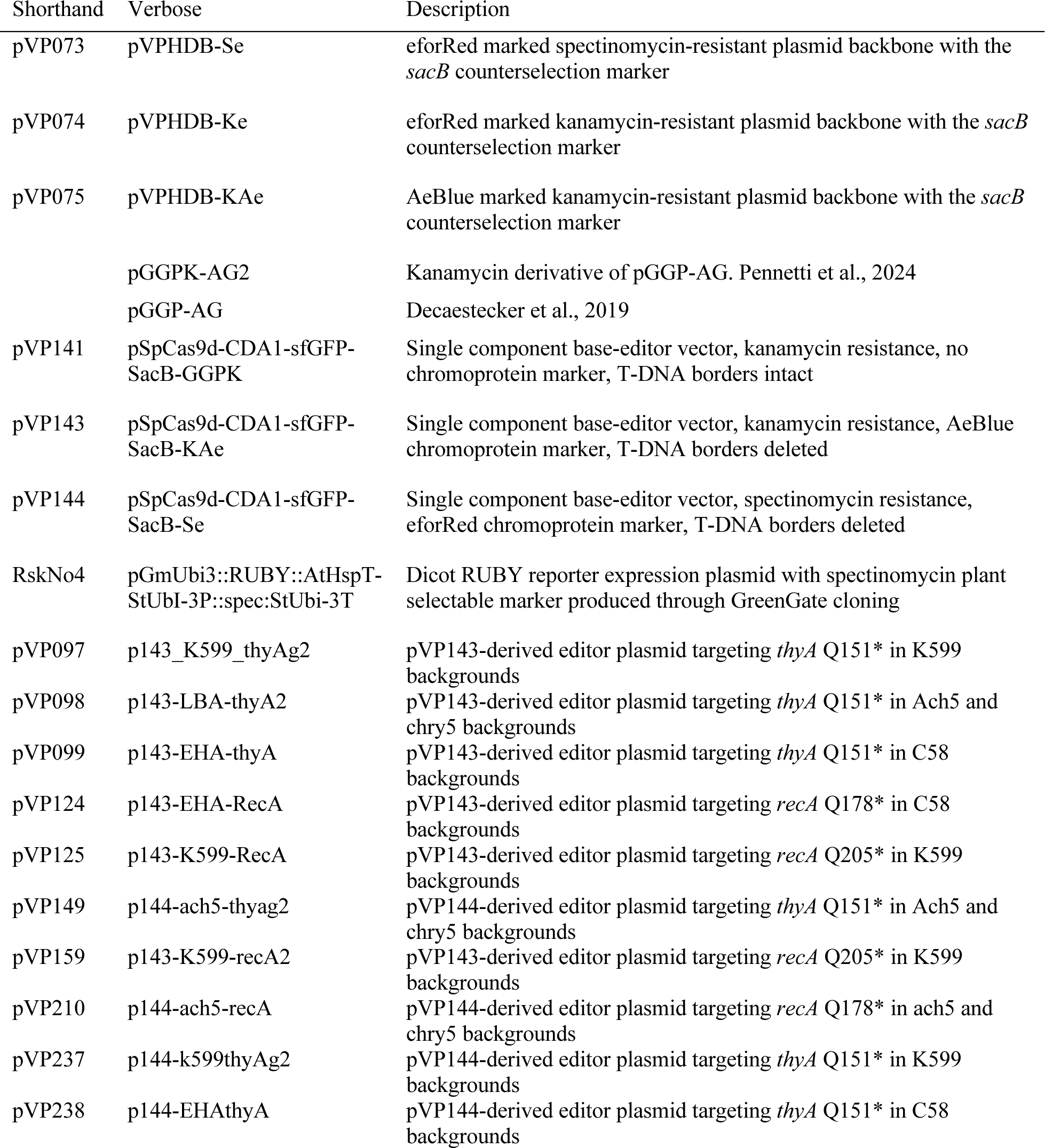
List of assemblies used in this study.

## Notes

### Competing Interest Statement

The authors have declared no competing interest.

